# Structural fuzziness of the RNA-organizing protein SERF1a determines a toxic gain-of-interaction

**DOI:** 10.1101/713511

**Authors:** N. Helge Meyer, Hanna Dellago, Carmen Tam-Amersdorfer, David A. Merle, Rosanna Parlato, Bernd Gesslbauer, Johannes Almer, Martha Gschwandtner, A. Leon, Titus M. Franzmann, Johannes Grillari, Andreas J. Kungl, Klaus Zangger, S. Fabio Falsone

## Abstract

Members of the MOAG/SERF protein family have been attributed a neuropathologic significance because of their ability to enhance the proteotoxic polymerization of amyloidogenic proteins such as alpha-Synuclein (aSyn). However, the cellular function remains unknown. Here, we identify SERF1a as an RNA-organizing protein i) with no structural homology to canonical RNA-binding proteins, ii) with an RNA-chaperoning activity which favours the incorporation of RNA into nucleoli and liquid-like artificial RNA-organelles, and iii) with a high degree of conformational disorder in the RNA-bound state. We demonstrate that this type of structural fuzziness determines an undifferentiated interaction of SERF1a with aSyn and RNA. Both molecules bind to one identical, positively charged site of SERF1a by an analogous electrostatic binding mode, with similar binding affinities, and without any observable disorder-to-order transition. The absence of primary or secondary structure discriminants results in SERF1a being unable to distinguish between nucleic acid and amyloidogenic protein, leading the pro-amyloid aSyn:SERF1a interaction to prevail under conditions of cellular stress. We suggest that structural fuzziness of SERF1a accounts for an adverse gain-of-interaction which favours toxic binding to aSyn at the expense of non-toxic RNA binding, therefore promoting a functionally distorted and pathogenic process. Our results provide a direct link between structural disorder, amyloid aggregation and the malfunctioning of RNA-associated cellular processes, three hallmarks of neurodegenerative diseases.

## INTRODUCTION

The Modifier of Aggregation-4/Small EDRK-rich Factor (MOAG4/SERF) protein family has emerged as an age-associated potentiator of amyloid polymerization (1,2), a type of reaction typical for a series of prevalent and incurable neurodegenerative disorders including Parkinson’s disease, Alzheimer’s disease, Huntington’s disease, and prion-related encephalopathies (3). The pro-amyloid mechanism has been studied in detail especially for the human homologue SERF1a and the amyloidogenic representative alpha-Synuclein (aSyn) (4,5), an intrinsically disordered protein (IDP) implicated in synaptic vesicle fusion (6). The interaction leads to a partial deprotection of amyloidogenic, aggregation-prone elements, and consequently to an increased propensity for amyloid polymerization. This process accelerates the generation of neuronal aSyn inclusions, a hallmark of synucleinopathies (7).

We have previously shown that the interaction between aSyn and SERF1a takes place in the cytosol to form a weak, short-lived, and dynamic complex (4). In their bound state, both proteins retain an unusually high degree of conformational disorder which is typical of random fuzzy complexes, i.e. of two IDPs interacting without stable folding (8). This type of molecular recognition is in sharp contradiction to the traditional structure-function paradigm which states that polypeptides underlie a strict folding hierarchy for a precise biological process.

Although native structural disorder allows for conformational adaptation, it also accounts for polypeptide misfolding and functional derangements. Considering the proteotoxic character of the highly flexible aSyn:SERF1a complex (1), it sounds reasonable to suppose that this interaction does not reflect a physiological function of SERF1a. The real functional properties of SERF1a in the cell, however, are still unknown.

Here, we report that structural fuzziness of SERF1a accounts for one physiologic and one pathogenic interaction deriving from undifferentiated binding to two separate species of molecules. We show that SERF1a is a dynamic RNA-binding protein (RBP) without any apparent structural homology to canonical globular RNA binding domains. We found that under physiologic conditions, SERF1a accumulated in the nucleus and more diffusely in the cytosol. In the nucleus, SERF1a localized to RNA-processing structures, clustering preferentially with networks of splicing and ribosomal RBPs especially including the nucleolus. By means of its RNA-chaperoning activity, SERF1a was able to mediate RNA-RNA interactions, accelerating the annealing of single complementary RNA strands, and facilitating their incorporation into nucleoli and artificial mimics of membraneless RNA-organelles. However, under stress conditions, SERF1a was rapidly released from the nucleolus, favouring amyloidogenic, and thus toxic binding to aSyn in the cytosol. The dual interaction with RNA and aSyn derived from a fuzzy binding mode, which was characterized by i) full disorder in complex, irrespective of the binding partner, ii) same-site binding to a linear stretch of basic residues, iii) analogous electrostatic interactions, and iv) similar K_D_-values for both complexes. Thus, protein fuzziness determines the inability of SERF1a to discriminate between the physiologic (RNA) and the pathogenic (aSyn) binding partner, so that cellular localization only seems the discriminating principle for binding partner decisions.

## RESULTS

### Nucleocytosolic distribution of SERF1a

To determine where and whether SERF1 is constitutively expressed, we performed an extensive analysis by immunofluorescence in readily available immortalised laboratory cell lines (HeLa ovarian carcinoma and SH-SY5Y neuroblastoma cells), primary tissue-derived cells (rat astrocytes in mixed glial preparations), and mouse brain sections (hippocampal neurons). SERF1a was present in the nucleus as well as in the cytosol (Fig. S1). In the cytosol, SERF1a was present either diffusely in HeLa, SH-SY5Y, or in single puncta in rat astrocytes. Intranuclear SERF1a was found in the nucleolus, Cajal bodies and nuclear gems (Fig. S2), as established by co-localizaiton with the specific marker proteins. The bipartite localisation pattern of endogenously expressed SERF1a coincided with that of transiently overexpressed SERF1a (ref and Fig. S1E).

Similarly to numerous nucleolar proteins (9), SERF1a was dynamically released from the nucleolus in response to established stressors of nucleolar homeostasis, such as heat, chemical/genetic transcriptional damage, and oxidative stress. A short time exposure of living cells to heat led to a redistribution of the SERF1a fluorescence signal from the nucleolus to the nucleoplasm and cytosol (Fig. 1A), suggesting a rapid transfer between these compartments. A similar effect was observable upon treatment with the generic oxidising agent H_2_O_2_ (Fig. 1B) or with ROS-inducing mitochondrial toxins 1-Methyl-4-phenyl-1,2,3,6-tetrahydropyridin (MPTP) (Fig. 1B) and rotenone (Fig. 1C), which collectively led to a visible reduction of the nucleolar SERF1a fluorescence signal. Similarly, nucleolar damage following the inhibition of rRNA synthesis either by the pharmacologic inhibitor actinomycin D in neuroblastoma cells (Fig. 1C), or by conditional genetic knockout of the nucleolar transcription factor TIF-IA in mouse brains, caused a loss of the nucleolar SERF1a fraction (Fig. 1D and supplementary results “*TIF-IA cKO mouse model”*). Expression levels of SERF1a were generally not decreased in response to stressors (Fig. S3), supporting a spatial rearrangement at protein level.

**FIGURE 1:**
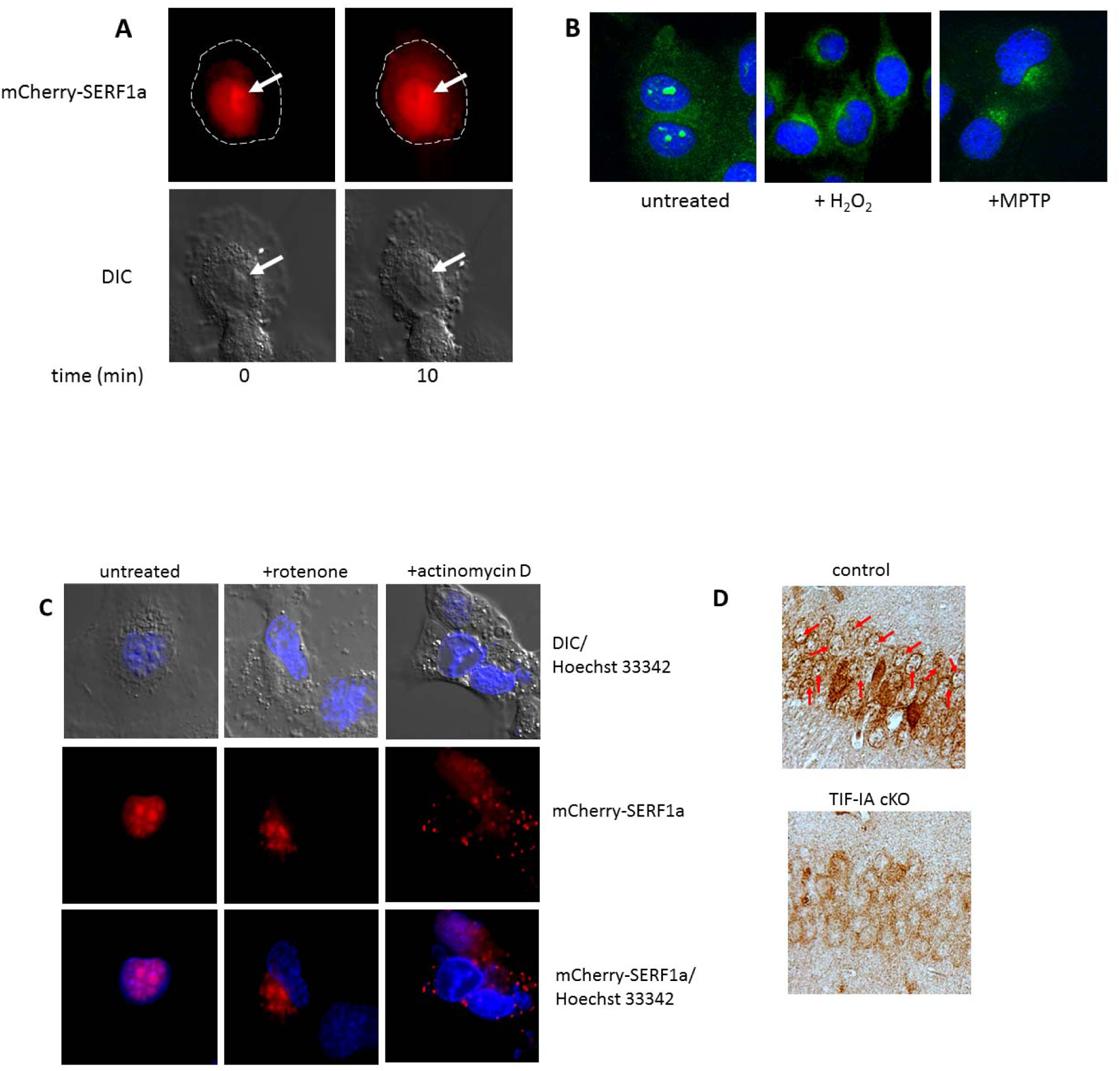
Stress-induced dynamics of SERF1a. (A) Upper panels: exposure of living SH-SY5Y cells to 42°C for 10 min causes the nucleocytosolic redistribution of overexpressed mCherry-SERF1a (red). Lower panels: corresponding DIC images. White arrows indicate the position of the nucleolus. (B) IF-staining of fixed SH-SY5Y cells treated with H_2_O_2_ or with MPTP. Endogenous SERF1a is green. Nuclei are stained with DAPI (blue). (C) SH-SY5Y living cells overexpressing mCherry-SERF1a (red) exposed to rotenone or actinomycin D. (D) Immunohistochemistry on paraffin sections of adult mouse brains showing SERF1a protein subcellular localization in hippocampal neurons of control (upper panel) and TIF-IA^CaMKCreERT2^ mutant (TIF-IA cKO) mice (lower panel) 4 weeks after induction of TIF-IA ablation by tamoxifen injection. Red arrows highlight the position of nucleoli. See also the section “*TIF-IA cKO mouse model”* in the supplementary results.

### SERF1a is an RNA-binding protein

To understand the functional relevance of SERF1a in the nucleus, a SERF1a-directed pull-down assay was performed out of nuclear extracts from cells overexpressing an enhanced green fluorescent protein-SERF1a (eGFP-SERF1a) chimera as bait (Fig. S4). The pull-down fraction was clearly enriched in RNA-binding proteins (RBPs) clustering to two major functional categories, namely ribosomal synthesis and RNA splicing (Fig. 2 and Table S1). This is in good correlation to the immunochemically determined localization of SERF1a in nucleoli (the sites of ribosome assembly), and in gems/Cajal bodies (the sites of spliceosome assembly). (Fig. S2). Accordingly, we found that SERF1a was able to influence i) splice site selection of an adenoviral minigene in living cells (see supplementary results “*Cellular splicing model*” and Fig. S5), and ii) expression levels of 47S pre-rRNA, a signature of ribosome biogenesis (ref) (see supplementary results “*Variations in pre-rRNA levels”* and Fig. S6).

**FIGURE 2:**
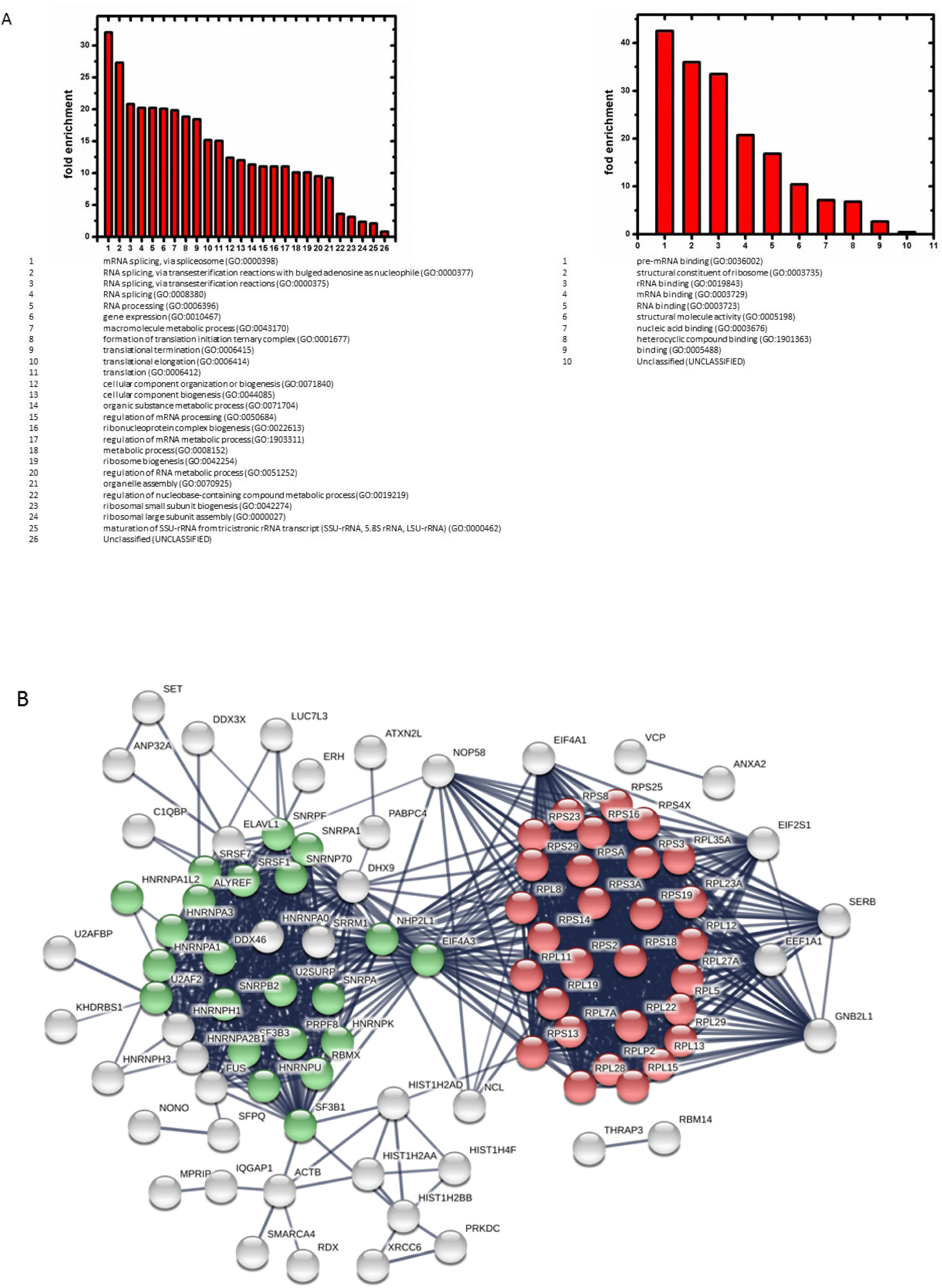
(A) PANTHER GO classification of proteins enriched out of nuclear extracts by SERF1a-directed pull-down, with respect to biological (left) and molecular (right) function. (B) STRING clustering analysis of the pull-down components. The network displays non-random clustering, with an average local clustering coefficient of 0.74, and a PPI enrichment p-value <10^−16^. Ribosome constituents are red. Spliceosome constituents are green. Disconnected nodes are not shown.

Due to the predominance of RBPs in pull-down fractions, we assumed the existence of a direct RNA:SERF1a interaction. This was supported by the protein’s basic net charge (pI=10.44) typical of nucleic acid binding proteins (Fig. 3A). SERF1a is absent from current comprehensive RBP catalogues (10), and similarity search algorithms failed to provide the existence of canonical RNA-binding domains. Interestingly, however, SERF1a shows low level homology to the RNA binding region of the splicing regulator matrin-3 (11,12). Moreover, by aligning SERF1a, which is intrinsically disordered, against a peptide database composed of 289 low complexity, non-canonical RNA-binding clusters (10), 2 potential RNA-binding regions were predicted (amino acids 7-20, and 49-54 (Fig. 3B)).

**FIGURE 3:**
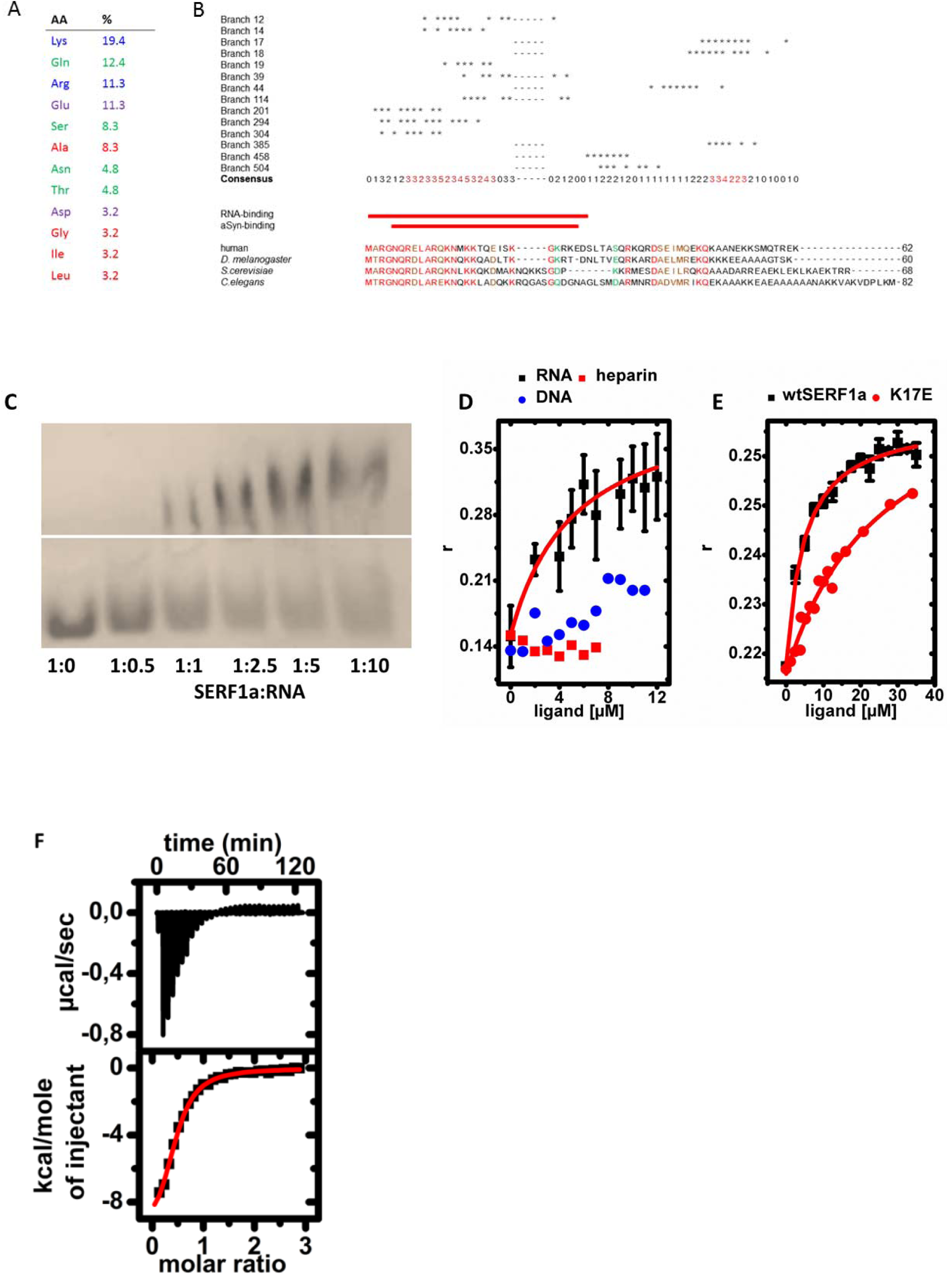
SERF1a binds directly to RNA. (A) Amino acid composition of SERF1a. (B) Putative RNA-binding regions derived by local similarity alignment of SERF1a against 289 intrinsically unstructured, non-canonical RNA binding motif clusters (branches) from the RBDmap database (ref). 14 branches were used for refinement. Numbers in red indicate SERF1a regions with highest consensus. Upper red bar: experimentally confirmed RNA-binding region (see also Fig. 6A). Lower red bar: aSyn binding region (ref). Sequence alignment shows the evolutionary conservation of the RNA-binding region. Identical amino acid residues are red. (C) Electrophoretic mobility shift assay (EMSA) of SERF1a with increasing concentrations of R21(-). (D) Fluorescence anisotropy saturation curves obtained by titration of Atto550-labeled SERF1a with R21(-) (■), heparin from porcine intestinal mucosa (■), or double stranded DNA 21mer (●). (E) Fluorescence anisotropy saturation curves obtained by titration of Cy3-R21(-) with wild-type SERF1a (■) or with point mutant K17E (●). (F) Isothermal titration calorimetry binding curve of the SERF1a:R21(-) interaction.

To test for direct RNA binding, we used purified recombinant SERF1a and a deliberately chosen, readily available synthetic RNA oligomer (R21(-)) for an electrophoretic mobility shift assay (EMSA). The addition of R21(-) to SERF1a caused significant concentration-dependent electrophoretic band shifting (Fig. 3C), corroborating a direct SERF1a:RNA interaction. We established the binding affinity of the complex by fluorescence anisotropy and isothermal titration calorimetry (ITC). As shown in Figures 3D and 3E, the addition of either R21(-) to Atto550-labeled SERF1a, or *vice versa* the addition of SERF1a to Cy3-labeled R21(-) led to saturable anisotropy binding curves with similar K_D_-values of 4.39±1.59 µM and 4.81±0.48 µM, respectively. ITC (Fig. 3F) yielded a comparable K_D_-value of 5.99±2.32 µM. Such values are indicative of a weak binding affinity typical of interactions with variable binding partners, as frequently found for highly flexible proteins (13). Consistent with the intrinsically disordered character of SERF1a, we found that this protein was capable of promiscuous binding to other types of RNA, such as tRNA (K_D_ = 1.10±0.76 µM) or total RNA from *S. cerevisiae* (K_D_=1.79±0.79 µM) (Fig. S7). However, it was unable to bind double stranded DNA or chemically unrelated polyanions such as heparin (Fig. 3D), indicating that binding to RNA was a selective process.

By ^1^H, ^15^N-HSQC NMR chemical shift titrations of isotope labelled SERF1a with 21R(-) (Fig. S8), we could pinpoint the RNA-binding region to an extended N-terminal stretch of mainly positively charged amino acids (Fig. 6A, upper panel), in correspondence to one of the computationally predicted RNA-binding regions (amino acids 6-28; Fig. 3B). From sequence alignments, we found that this site displayed the highest similarity among MOAG-4/SERF family members (Fig. 3B). We could indeed measure similar K_D_-values of 4.79±1.34 µM and 2.35±0.56 µM, respectively, for the eukaryotic SERF1a orthologues MOAG-4 from *C. elegans* (42% identity) or YDL085C-A from *S. cerevisiae* (44% identity) (Fig. S7), which suggests that RNA binding is evolutionary conserved. The functional relevance of the RNA-binding region was accentuated by a high charge sensitivity, as the replacement of one single conserved charged residue, e.g. an arginine to glutamate substitution at position 17 (K17E) was sufficient to significantly alter the K_D_-value of the complex from approx. 4µM to >100 µM (Fig. 3E). Collectively, these findings clearly identify SERF1a as an RBP.

We used fluorescence-labelled recombinant SERF1a together with purified nuclei and nucleoli to test *in vitro* whether RNA binding was necessary for the spatial organization of SERF1a in the cell. Firstly, Atto647N-SERF1a, when directly applied to isolated nuclei, was able to diffuse passively into the nucleoplasm, readily accumulating inside nucleoli (Fig. 4A). In contrast, the RNA binding-defective SERF1a point mutant Atto647N-K17E, while still able to diffuse into nuclei, was not enriched in nucleoli. Secondly, Atto647N-SERF1a clearly localized to isolated nucleoli (Fig. 4B), whereas Atto647N-K17E remained diffuse in solution. Thirdly, overexpressed eGFP-SERF1a was enriched in the nucleolus of HeLa living cells (Fig. 4C), while eGFP-K17E was excluded from the nucleolus. These results indicate that RNA binding defects can compromise nuclear partitioning of SERF1a. Thus, the nucleolar localization of SERF1a derives from a direct interaction with RNA.

**FIGURE 4:**
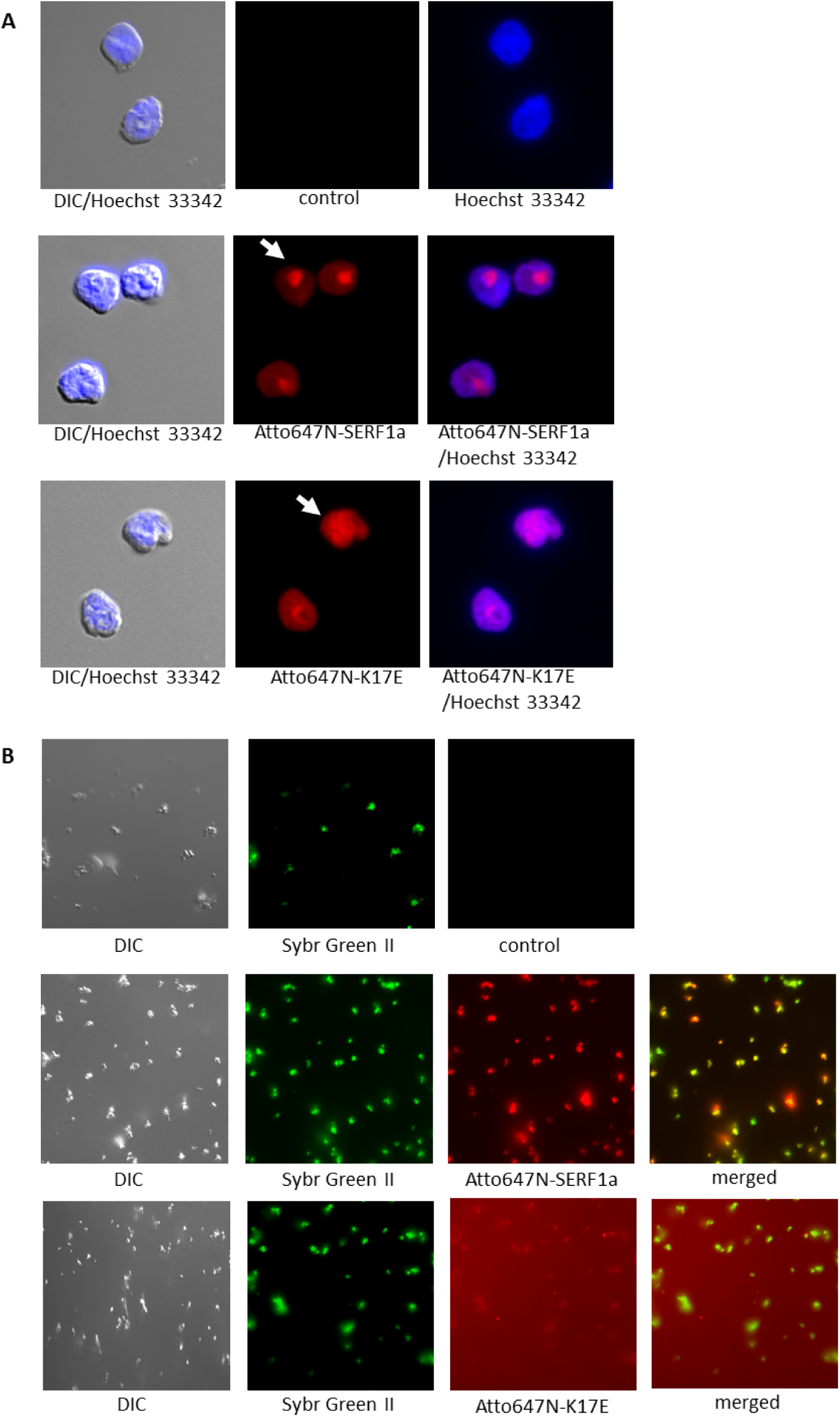
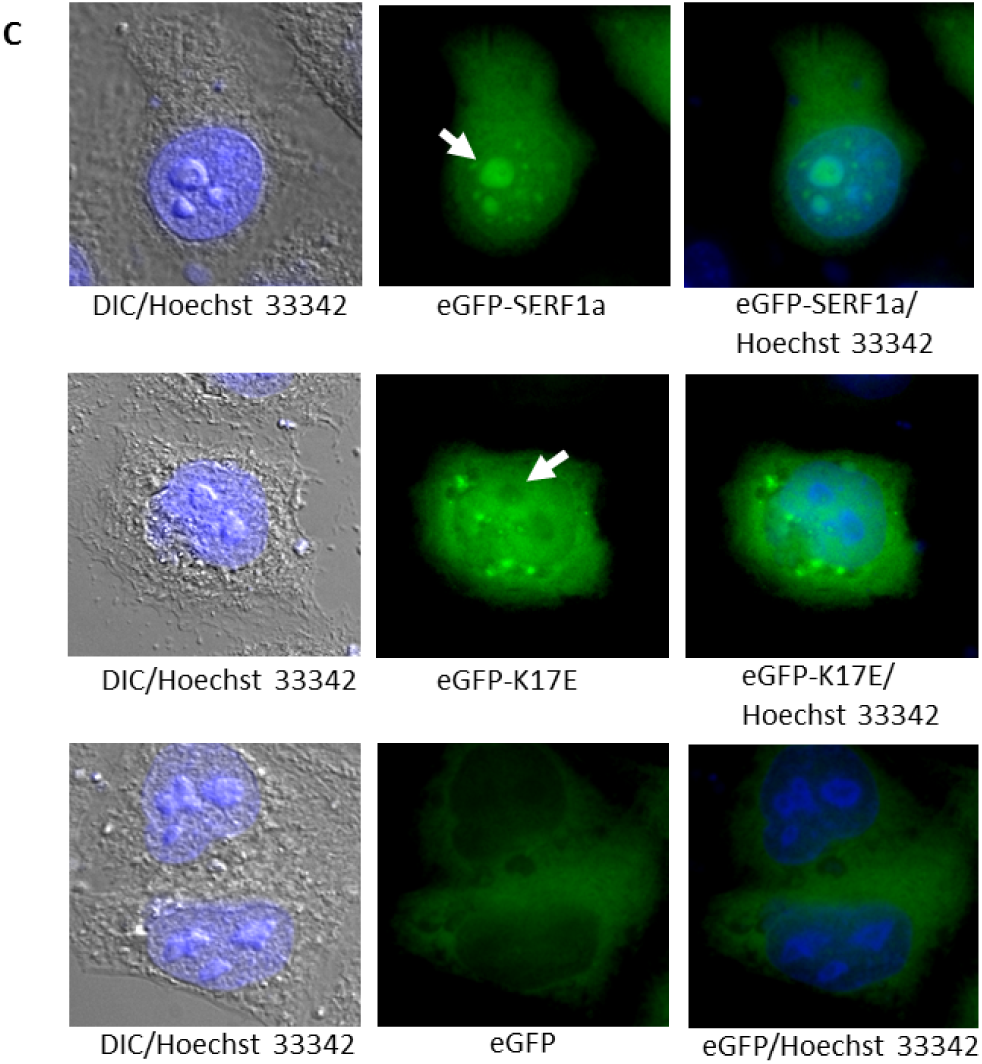
RNA binding determines localization of SERF1a. (A) Fluorescence imaging of isolated untreated nuclei (upper panel), of nuclei incubated with 10µM Atto647N-SERF1a (middle panel), and of nuclei incubated with 10 µM Atto647N-K17E. Arrows indicate the nucleolar accumulation of fluorescent wildtype SERF1a in the nucleoli, and the absence of the RNA-binding defective SERF1a mutant K17E from nucleoli. Nuclei were counterstained with Hoechst 33342 (blue). (B) Fluorescence imaging of untreated isolated nucleoli (upper panel), of nucleoli incubated with 10µM Atto647N-SERF1a (middle panel), and of nuclei incubated with 10 µM Atto647N-K17E. Nucleoli were counterstained with Sybr Green II RNA stain (green). (C) Overexpression of eGFP-SERF1a (upper panel) of the RNA-binding defective SERF1a mutant eGFP-K17E (middle panel), and of eGFP control vehicle (lower panel) in HeLa cells. Arrows indicate the accumulation of fluorescent wildtype SERF1a in nucleoli, and the exclusion of K17E from nucleoli.

### SERF1a is an RNA-organizing protein

Structural disorder in RBPs has been associated with the ability to favour functionally correct base pairing (14). With this respect, we tested SERF1a for a so-called RNA-chaperone activity in an *in vitro* based RNA-annealing fluorescence resonance energy transfer (FRET) assay (15). *Trans*-annealing of two complementary RNA 21mers, labelled with a FRET donor or acceptor dye respectively (cy5-R21(+) and cy3-R21(-)), leads to an increase of the FRET signal, which can be used to derive kinetic parameters of the reaction (15). Purified SERF1a remarkably improved RNA duplex formation (Fig. 5A), causing an increase of the annealing reaction constant K_obs_ by one order of magnitude from 0.0056±0.00026 s^−1^ (lit. 0.005 s^−1^ (15)) to 0.062±0.0021 s^1^. A significantly minor increase in K_obs_ (0.033±0.0015 s^−1^) was observed for the RNA-binding defective SERF1a mutant K17E. These results identify SERF1a as being able of promoting RNA-RNA base pairing.

**FIGURE 5:**
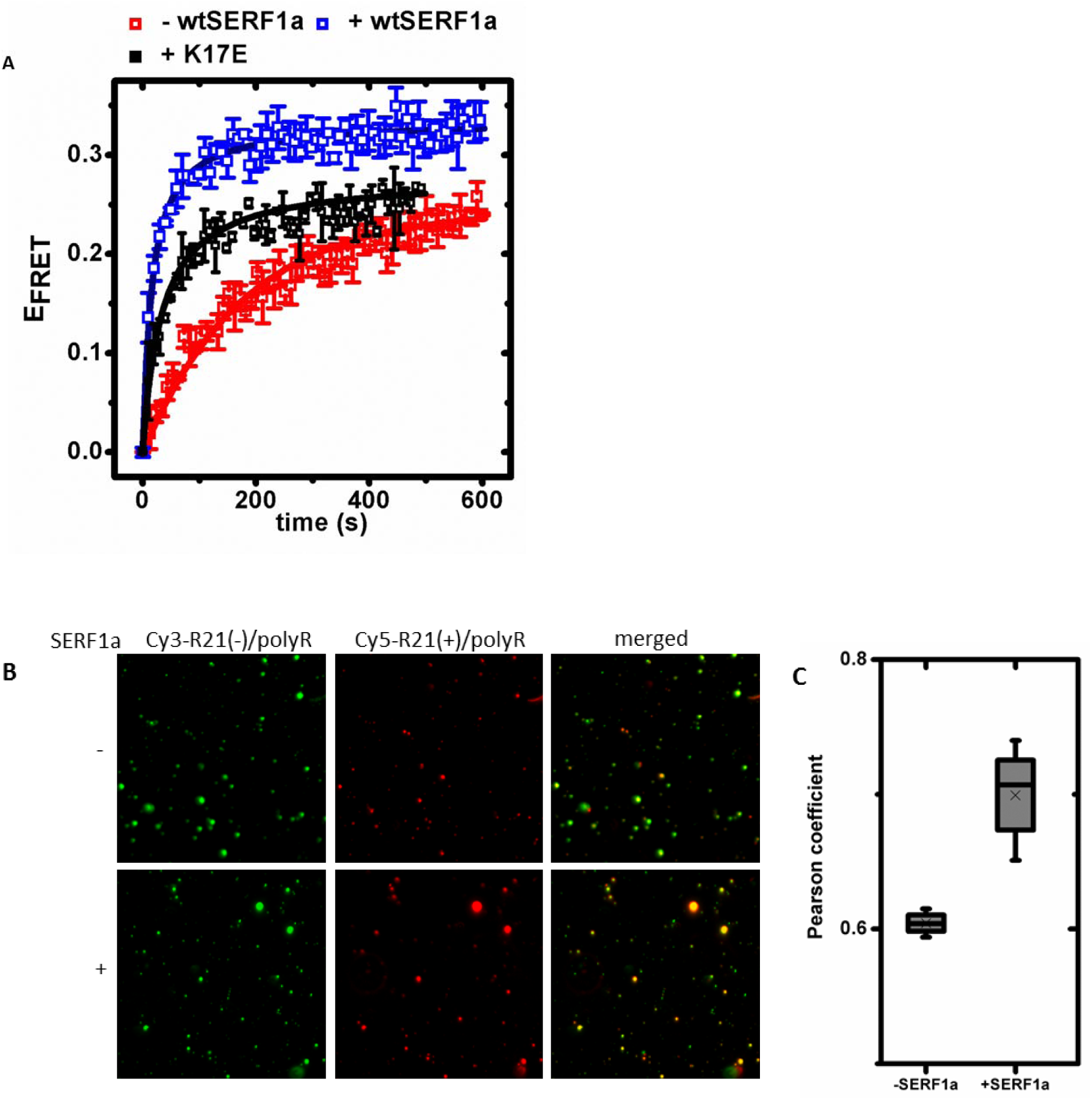
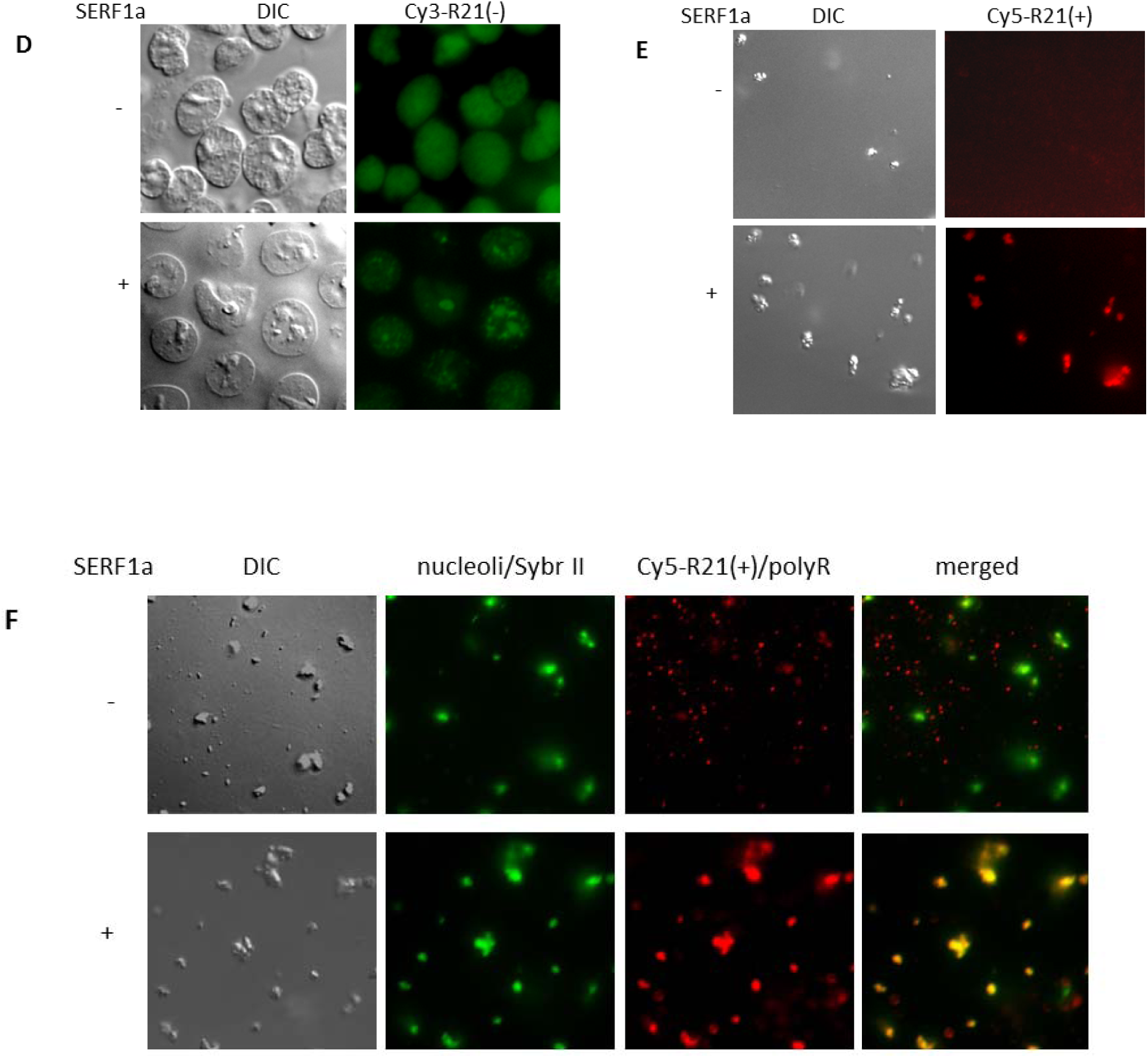
SERF1a is an RNA chaperone. (A) Kinetics of RNA complementary strand annealing in the absence (■) and in the presence of SERF1a(■) or of the RNA-binding defective SERF1a point mutant K17E (■). (B) Co-localization between Cy3-21R(-)/polyR and Cy5-21R(+)/polyR complementary complex coacervates in the absence (-) and in the presence (+) of 10 µM SERF1a. (C) Co-localization between between Cy3-21R(-)/polyR and Cy5-21R(+)/polyR in the absence and in the presence of 10 µM SERF1a, expressed as Pearson coefficient. Crosses indicate mean values of three independent experiments. (D) Diffusion and partitioning of Cy3-21R(-) into isolated nuclei in the absence (-) and in the presence (+) of 10 µM SERF1a. (E) Incorporation of Cy5-21R(+) into isolated nucleoli in the absence (-) and in the presence (+) of 10 µM SERF1a. (F) Incorporation of Cy5-21R(+)/polyR complex coacervates into isolated nucleoli in the absence (-) and in the presence (+) of 10 µM SERF1a. Nucleoli are counterstained with Sybr Green II RNA stain (green).

RNA-RNA interactions contribute to the structural organization of RNA-rich membraneless organelles (16–21). *In vitro*, basic physical properties of membraneless biological compartments can be mimicked by RNA-derived liquid droplets, which are useful artificial models for studying fluidity, fusion behaviour, and composition dynamics (18–21). We generated fluorescent RNA-based complex coacervates, which form by liquid-liquid demixing of oppositely charged polymers (21), to analyse SERF1a-mediated RNA-RNA annealing at two phase regime, i.e. under conditions where RNA is present as a phase separated condensate (Fig. 5B). The addition of polyarginine to Cy3-R21(-) led to a spontaneous formation of fluorescent liquid droplets with a diameter of approximatively 1 µm. Fluorescence signal co-localization by the addition of complementary Cy5-R21(+)/polyR droplets was an indication for strand annealing in the coacervate phase. In the presence of SERF1a, co-localization of Cy5-R21(+)/polyR and Cy3-R21(-) was increased, as deduced by a change from 0.604 to 0.699 in the Pearson correlation coefficient (Fig. 5C), which is one quantitative measure of co-localization (22). Thus, the incorporation of complementary RNA into the coacervate was facilitated by the RNA-chaperone properties of SERF1a.

We assumed by conceptual analogy that SERF1a might chaperone the incorporation of RNA into nucleoli, as these organelles share physical properties of liquid droplets and are highly enriched in RNA (23, 24). In a simplified approach, we directly added RNA to intact nuclei or to isolated nucleoli. The fluorescent probe Cy3-R21(-) was able to enter the nuclei by diffusion, randomly distributing inside the nucleoplasm (Fig. 5C). Upon the simultaneous addition of purified SERF1a, we noticed local repartitioning of the fluorescence signal into non-random patterns suggestive of subnuclear compartments. The corresponding DIC images showed that prominent signals derived from nucleoli and less prominent subnuclear particles. Similarly, the probe Cy5-R21(+) failed to target isolated nucleoli as a monomer (Fig. 5D) or as a phase separated coacervate (Fig. 5E) in the absence of SERF1a, while becoming visibly enriched into nucleoli in the presence of SERF1a. We deduced that SERF1a favoured the nucleolar integration of RNA. Collectively, these results suggest an RNA-organizing role of SERF1a inside the nucleus.

### RNA and aSyn interact with SERF1a by a similar binding mode

Next, we aimed to understand how SERF1a, which is a known modifier of amyloid protein aggregation (1,2), distinguishes between RNA and amyloidogenic protein partners such as aSyn, and consequently whether both processes can influence each other. We initially hypothesized the existence of two distinct binding regions for the separate recognition of RNA and amyloidogenic substrate. However, NMR-derived RNA:SERF1a and aSyn:SERF1a interaction data revealed a remarkable coincidence of the two binding interfaces. As shown by comparison in Fig. 6A, prominent ^1^H, ^15^N-HSQC NMR chemical shift perturbations caused by the addition of aSyn or 21R(-) to ^15^N-labeled SERF1a invariably occurred within one positively charged, linear amino acid motif (approximatively residues 1-30) at the N-terminus of SERF1a (see also Fig 3B). These findings contradict the existence of two distinct binding sites.

**FIGURE 6:**
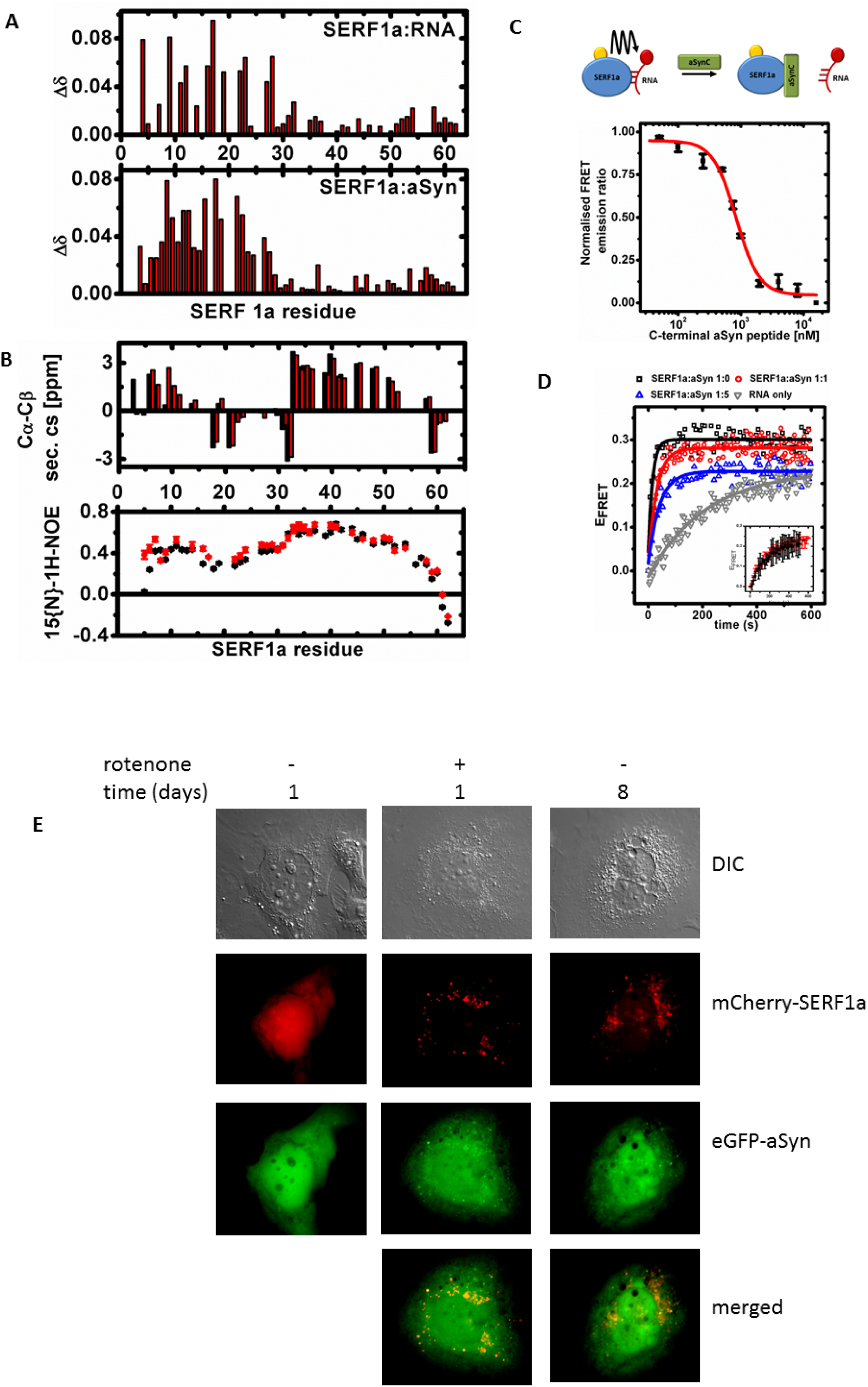
SERF1a binds promiscuously to RNA and aSyn. (A) Chemical shift changes Δδ of ^15^N-SERF1a amino acid residues upon the addition of R21(-) (upper panel) or aSyn (lower panel; adapted from Merle et al. (4)). (B) Upper panel: 13C^α/β^ secondary chemical shifts of SERF1a either free (black) or in complex with R21(-) (red); lower panel: {^1^H}-^15^N heteronuclear NOE values of SERF1a free (●) or in complex with R21(-) (♦). (C) Upper scheme: experimental setup of FRET-based competition assay. The displacement of RNA from SERF1a by aSyn was measured by FRET signal changes of a pre-formed Atto550-SERF1a:Cy5-R21(+) donor-acceptor complex upon the addition of an unlabeled C-terminal aSyn peptide (aSynC; aa 100-140). Lower panel: FRET competition curve obtained from fluorescence signal changes of the Atto550-SERF1a:Cy5-R21(+) complex upon titration with aSynC. (D) FRET-based RNA duplex annealing kinetics of Cy3-R21(-) and Cy5-R21(+) in the absence (∇), and in the presence of SERF1a (□), and of SERF1a with increasing amounts of aSyn (SERF1a:aSyn 1:1, ○; SERF1a:aSyn 1:5, Δ). Control insert: RNA duplex annealing kinetics in the absence (red) and in the presence of aSyn (black). (E) SH-SY5Y living cells overexpressing mCherry-SERF1a (red) or eGFP-aSyn (green) 1 day or 8 days after transfection in the absence or in the presence of 200 nM rotenone.

We hypothesized next that SERF1a might distinguish between RNA and aSyn by secondary structure. We analyzed if RNA induces stable folding of intrinsically disordered SERF1a as opposed to aSyn, which does not induce secondary structure upon binding (4). However, this hypothesis was disproved by (A) an invariably narrow ^1^H, ^15^N-HSQC NMR proton signal dispersion (8.0-8.8 ppm), indicative of structural disorder of SERF1a in the free and RNA-bound state (25) (Fig. S8), (B) little ^13^C^α^ and ^13^C^β^ secondary chemical shift (^13^C^α/β^-SCS) signal variation, reflecting an unaffected folding propensity of the SERF1a backbone upon the addition of RNA (26) (Fig. 3B, upper panel), and (C) {^1^H}-^15^N heteronuclear Nuclear Overhauser Effects (NOEs) of RNA-bound SERF1a remaining generally below those expected for rigidly folded structure (around 0.8) (27) (Fig. 3B, lower panel). Based on these results, we concluded that SERF1a remains structurally disordered in complex irrespectively of RNA or aSyn as a binding partner.

Collectively, these findings demonstrate that SERF1a is unable to distinguish between nucleic acid or protein binding by means of primary or secondary structure, which was reflected by comparable binding affinites of the aSyn:SERF1a and the RNA:SERF1a interaction (K_D_-values between 4-7 µM (2)). Consequently, we supposed that aSyn and RNA binding might interfere with each other. We thus established a FRET *in vitro* competition assay based on the addition of an unlabelled C-terminal aSyn peptide (aSynC) corresponding to the SERF1a binding site of aSyn (2,4), to a preformed Atto550-SERF1a:Cy5-R21(-) FRET donor:acceptor complex (Fig. 6C). By increasing the amounts of aSynC, we observed a concentration-dependent sigmoidal decrease of the energy transfer signal, indicating that an excess of aSyn could efficiently displace RNA from SERF1a (IC_50_ = 840±64 nM), thereby disrupting FRET. In line with this observation, the RNA-annealing efficiency of SERF1a gradually decreased by increasing aSyn concentrations (Fig. 6D). Because free aSyn did not bind RNA (Fig. S9), neither did it directly influence RNA annealing (Fig. 6D, control insert), we deduced that the FRET signal change derived from a direct competition of aSyn for SERF1a binding.

Due to this concurrent binding mode, we supposed that the local cellular abundance of the interacting partners might determine the prevalence of one particular interaction. Considering the toxic character of the purely cytosolic aSyn:SERF1a interaction, we investigated in particular the effects of increased cytosolic SERF1a levels upon nucleolar stress. Doubly transfected SH-SY5Y cells co-overexpressing eGFP-aSyn and mCherry-SERF1a were subjected to intoxication with rotenone, a suspected inducer of Parkinson’s disease (28) which triggers a nearly quantitative transfer of SERF1a from the nucleus to the cytosol (Fig. 1C). The aSyn:SERF1a interaction is characterized by the time-dependent appearance of granular cytosolic bodies incorporating both proteins (4). In untreated cells, this process became typically observable 5-8 days after transfection (Ref. 4 and Fig. 6E, right panel). In rotenone-treated cells, it was considerably accelerated by the forced release of SERF1a into the cytosol, being observable one day after exposure (Fig. 6E, middle panel). These results suggest that an increase of cytosolic SERF1a concentrations, e.g. by nucleolar stressors, favours the highly amyloidogenic interaction of SERF1a with aSyn in the cytosol.

## DISCUSSION

In this study, we identified SERF1a as a highly disordered RBP with no apparent structural homology to canonical RNA-binding elements. SERF1a was able to bind different types of RNA molecules, suggesting an unspecific binding mode which was reflected by a low substrate affinity, a generally unspecific RNA-chaperone activity, and clustering to multiple types of RNA-associated protein networks. SERF1a showed binding selectivity, as it distinguished single stranded from double stranded nucleic acids or chemically unrelated polyanions. A stable conformation was not required for selectivity, as the interaction occurred in the absence of folding, which is typically referred to as fuzzy binding (8). Instead, RNA binding, function and spatial distribution of SERF1a was uniquely determined by a highly conserved primary structure charge composition, and minimal charge variation significantly decreased the affinity for RNA, disrupted nucleolar localization, and affected SERF1a function.

Our results demonstrate previously unreported same-site binding of a nucleic acid and a polypeptide to a fully disordered protein, accounting for an undifferentiated, highly disordered binding mechanism characterized by comparably weak binding affinities. Intrinsic structural disorder is a regulatory feature of RBPs, as systematically shown by the existence of hundreds of structureless RNA binding motifs counterposed to traditional globular RNA-binding domains (28–30). However, conformational flexibility in protein complexes also increases the risk of misfolding. We suppose that structural fuzziness of SERF1a, while of advantage for promiscuous RNA interactions, increases the tendency of binding malfunctions. Indeed, it was previously shown that SERF1a is also responsible for the amyloidogenic rearrangement and toxic aggregation of aSyn in consequence to a fully disordered interaction (2,4). The indiscriminate binding of aSyn and RNA to SERF1a *via* one binding site points to an interference which favours the conversion of SERF1a from an RNA-organizing protein into a neurotoxic pro-amyloid factor, with cellular localization as the sole discriminating principle of binding partner decisions. This is intriguing, considering that a) RNA pathways are frequently compromised in neurodegenerative disorders (31,32,33), and b) several RBPs enriched by SERF1a pull-down capture are capable of amyloid polymerization (34). Due to similar K_D_-values of both complexes, it is reasonable to suppose that, in the cell, one complex can prevail depending on the local abundance of RNA, aSyn and/or SERF1a levels. Under non-stress conditions, SERF1a resides predominantly in the nucleolus, which is highly enriched in RNA (up to 40 mg/ml) (35). Although mutant aSyn can be expressed in the nucleolus in transgenic mice (36), wild type aSyn is typically absent from the nucleolus, which prevents competition with RNA. Our results show that the stress-induced expulsion of SERF1a into the cytosol favours binding to aSyn, an interaction which might be forced by persistent nucleolar stress. The interaction with aSyn might be further facilitated by cytosolic RNA levels which are well below those of the nucleus (37). Finally, this interaction might be forced by a pathogenic excess in aSyn levels, e.g. by PD-associated *SNCA* gene number variation (38) or by a defective homeostasis (39). As a general consequence, SERF1a-associated RNA pathways might deteriorate in favour of an amyloidogenic gain-of-interaction.

## MATERIALS AND METHODS

Animal experiments carried out by C.T. were in accordance with the guidelines and regulation of the Austrian Ministry of Research, Science and Economics. All experimental protocols were approved by the Austrian Ministry for Research, Science and Economics and by the local facility for animal experiments, Medical University of Graz, Institute for Biomedical Research, Graz, Austria. Animal experiments carried out by R.P. were approved by the Committee on Animal Care and Use (Regierungspräsidium Karlsruhe, 35-9185.81/G-180/08) in accordance with the local Animal Welfare Act and the European Communities Council Directives (2010/63/EU and 2012/707/EU).

Unless otherwise specified, all reagents were obtained from Sigma-Aldrich, Vienna, Austria. RNA strands 21R(+) (5′-AUGUGGAAAAUCUCUAGCAGU-3′), 21R(-) (5′-ACUGCUAGAGAUUUUCCACAU-3′), Cy5-21R(+) and Cy3-21R(-), and the two corresponding DNA strands were purchased from VBC-Biotech, Vienna, Austria. dsDNA was generated by cooling of a 1:1 molar mixture of single stranded DNA after heating at 95°C. aSyn C-terminal peptide GILEDMPVDPDNEAYEMPSEEGYQDYEPEA (aSynC, aa 100-140) was synthesized by Pepnome Inc (Hong Kong, China). HeLa cells were obtained from BioBank Graz, Medical University Graz, Austria. Undifferentiated SH-SY5Y neuroblastoma cells were from LGC Standards, Wesel, Germany.

### Mutagenesis

The *eGFP-K17E* construct for the mammalian expression of the SERF1a single-point mutant eGFP-K17E was generated by site-directed mutagenesis according to the QuickChange II mutagenesis kit manual (Agilent, Santa Clara, CA), using *pEGFP/SERF1a* (Tab. S3) as a mutagenesis template.

### Protein purification

The production of unlabeled and isotope labeled SERF1a, K17E, and aSyn has been described previously (2,4). MOAG-4 was a gift from E.A.A. Nollen, Groningen, The Netherlands. For the expression of YDL085C-A in *E. coli*, we used a sequence optimised synthetic gene cloned into the T7-inducible pJ414 vector (ATUM, Newark, CA). The protein was expressed and purified analogously to the human isoform SERF1a as described in Ref. 2.

The site-selective conjugation of SERF1a and K17E with Atto647N or Atto550 has been described in Ref. 2. See Tab. S3 for a comprehensive list of used plasmids.

### Antibodies

Polyclonal goat anti-human SERF1a (clone K-13), monoclonal mouse anti-human fibrillarin (clone B-1), monoclonal mouse anti-human SMN (clone F-5), monoclonal mouse anti-human Coilin (clone 56), and donkey anti-goat IgG FITC were purchased from Santa Cruz Biotechnology, Inc., TX, USA. Rabbit anti-mouse IgG (H+L) Cy3 was from Jackson ImmunoResearch Lab, Inc., West Grove, PA, USA. For IF, all antibodies were diluted with Dako Cytomation Antibody Diluent with background reducing components (Dako, Inc., Carpinteria, CA, USA).

### Induction of nucleolar stress

For living cell fluorescence microscopy, undifferentiated SH-SY5Y cells were grown and transfected with mCherry-SERF1a as described in the section *“Living cell microscopy”*. One day after transfection, cells were exposed to heat (42°C for 10 min.), rotenone (200 nM for 12 h), or actinomycin D (2µM for 3 h). For co-localization with aSyn, cells were co-transfected with mCherry-SERF1a and eGFP-aSyn. One day after transfection, cells were treated with 200 nM rotenone.

For IF microscopy, undifferentiated SH-SY5Y cells growing on SuperFrost slides were treated with H_2_O_2_ (500 µM for 30 min.) or MPTP (Axon Medchem, Groningen, The Netherlands) (100 µM for 12 h). Cells were further processed as described in the section below.

### Mouse treatment by tamoxifen

Homozygous TIF-IA^flox/flox^; CaMKCreERT2 (TIF-IA^CaMKCreERT2^) mutant mice were generated by crossing mice carrying the TIF-IA floxed allele (TIF-IA^flox/flox^) to the transgenic line CaMKCreERT2. TIF-IA^+/flox^; CaMKCreERT2 positive mice were again crossed with TIF-IA^flox/flox^ mice (40). 2-month-old TIF-IA^flox/flox^ control and TIF-IA^CaMKCreERT2^ mutant mice were both injected intraperitoneally with 1 mg tamoxifen twice a day for five consecutive days and were analyzed 4 weeks after the last injection as previously described (41).

### Immunofluorescence (IF) staining

Mouse brain isolation and IF staining of brain sections was performed as previously described (4). In brief, studies were carried out on 4% PFA fixed, paraffin embedded 10 days old C57BL/6 mouse brains. Polyclonal goat anti-human SERF1a (K-13) IgG and donkey anti-goat FITC (green) were used. Nuclei were counterstained with DAPI (Life Technologies/Thermo Fisher Scientific, Waltham, MA). Images were acquired on a Leica DM4000 B microscope equipped with Leica DFC 320 Video camera (Leica Cambridge Ltd).

Rat astrocytes were cultured as described previously (42). Primary cultures of glial cells were prepared from the brains of 2-day-old Wistar rats. After transfer of single cell suspension to culture flasks (1 brain/75 cm^2^) cells were cultivated for 6–8 days in DMEM supple-mented with 10% heat-inactivated fetal calf serum (FCS), 50 U/ml penicillin, and 50mg/ml streptomycin.Astrocytes were trypsinized from culture vessels and reseeded on 10 cm culture dishes containing poly-lysine-coated glass cover slides. Cells were cultivated for 7 days in DMEM supplemented with 10% FCS. Cover-slides were fixed with 4% PFA and stained with goat anti SERF1a/donkey anti goat FITC (green) and mouse anti glial fibrillar acidic protein/ anti-mouse Dylight 594 (red). Cover slides where mounted with Vectashield mounting medium (Vector Laboratories Inc., Burlingame, CA, USA) containing DAPI as nuclear counterstain.

IF staining of HeLa or SH-SY5Y cells was performed on cells cultured on SuperFrost slides (Thermo Fisher Scientific). Cells were fixed with cold acetone for 5 min. After fixation, cells were washed with Tris-Buffered Saline Tween-20 (TBST, pH7.4) and blocked with UV ultra-block (Thermo Fisher Scientific) for 7’ before primary antibodies incubation at 4°C overnight (anti SERF1a at 2.67µg/ml; anti SERF1a at 2.67 µg/ml-anti FBL at 0.33µg/ml; anti SERF1a at 2.67 µg/ml-anti SMN at 0.167µg/ml; anti SERF1a at 2.0 µg/ml-anti Coilin at 0.083 µg/ml). All incubation steps were performed in a dark moist chamber at room temperature. After 5 min. of TBST wash secondary antibodies were applied: Donkey anti-goat FITC (green) 0.50µg/ml; or donkey anti-goat FITC (green) 0.50µg/ml-rabbit anti-mouse Cy3 (red) 1.75µg/ml for 30 min. After rinsing in TBST, DAPI (blue) 5µg/ml was added to the slides for 20 min. as a nuclei counterstain. Cells were rinsed again with TBST before mounting with Vectashield mounting medium (Vector Laboratories Inc.).

### Histological analysis of hippocampal mouse tissue

Wild type and TIF-IA^CaMKCreERT2^ mutant mice were sacrificed by cervical dislocation and brains were immediately dissected. For immunohistochemistry, one brain hemisphere was fixed in 4% paraformaldehyde overnight and paraffin embedded, paraffin sections were 7 µm thick. Visualization of antigen-bound primary polyclonal goat anti-human SERF1a (K-13) (1:100) antibody on paraffin sections of adult mouse brains following antigen retrieval (HK086-9K, Biogenex, San Ramon, CA) was carried out using a biotinylated secondary antibody together with the avidin-biotin system and the VECTOR peroxidase kit (PK-6100, Vector Laboratories) using diaminobenzidine tablets as a substrate.

### Living cell microscopy

SH-SY5Y and HeLa cells were grown, respectively, in DMEM or DMEM/F-12 HAM medium supplemented with 10% FCS and penicillin/streptomycin antibiotics (PAA, Pasching, Austria) at 37□°C and 5% CO_2_. Cells were grown on 35□mm poly-D-lysine coated glass bottom dishes (MatTek, Ashland, MA). Unsilenced or previosuly silenced cells (see section “*Transient SERF1a silencing”*) were transiently (co)-transfected with the desired mammalian expression construct(s) (Tab. S3) using SuperFect transfection reagent (Qiagen, Hilden, Germany) according to the manufacturer’s manual. Differential intereference contrast (DIC) and fluorescence microscopy images were recorded at 37°C on a Zeiss Axio Observer Z1 inverted microscope (Zeiss, Jena, Germany) equipped with epifluorescence illuminator and plate heating chamber. Images were processed using the ImageJ software.

### Transient SERF1a silencing

SH-SY5Y cells were grown to semi-confluence at 37° C and 5% CO_2_ in Dulbecco’s Modified Eagles Medium supplemented with 10% foetal calf serum and antibiotics. Before silencing, cells were washed with PBS. The silencing reaction was performed in Gibco OPTIMEM medium (Thermo Fisher Scientific) by the addition of a 50 nM pre-designed siRNA primer pair CCAGGAAAUUAGCAAGGGATT (sense) and UCCCUUGCUAAUUUCCUGGGT (antisense) (Ambion/Thermo Fisher Scientific; siRNA ID# s15794) previously co-incubated for 20 min. with Lipofectamine (Thermo Fisher Scientific). After 24h, cells were washed with PBS and prepared for further processing (Fig. S10).

### qPCR

Total RNA from cellular samples was isolated using the GenElute Mammalian Total RNA Miniprep Kit RTN70. cDNA was produced by the High Capacity cDNA Reverse Transcription Kit (Thermo Fisher Scientific) according to the manufacturer’s protocol. qPCR was run on a an Applied Biosystems 7300 Real Time PCR System.

Total RNA from hippocampal vibratome sections (300μm) was extracted using the RNeasy Mini Kit (Qiagen). Reverse transcription was performed using random hexamers and MultiScribe Reverse Transcriptase (Applied Biosystems) following the manufacturer’s instructions. qPCR was run on a Light Cycler 480 (Roche, Basel, Switzerland). For each amplicon, serial dilutions of cDNA from the hippocampus were included in each run to generate standard curves for relative quantification by the instrument’s software.

The Power SYBR Green PCR MasterMix (Thermo Fisher Scientific) was used according to the manufacturer’s protocol. Used primers are listed in Table S2. The thermo cycle program consisted of an initial denaturation step (10 min. at 95 °C), followed by 40 cycles (45 cycles for 5’ ETS primer amplification) of denaturation (15 sec. at 95 °C), primer annealing (30 sec. at 60 °C) and elongation (1 min. at 72 °C). At the end, one final dissociation step (system default) was added.

### Isolation of nuclei and nucleoli, and localization analysis

Nuclei were purified using the Active Motif Nuclear Complex Co-IP Kit (Active Motif, Carlsbad, CA) as described in the manufacturer’s manual (www.activemotif.com/catalog/25/nuclear-complex-co-ip-kit). Briefly, 5⋅10^6^ SH-SY5Y cells were resuspended in hypotonic buffer, incubated for 15 min. on ice, and lysed by the addition of 1% kit detergent. Nuclei were collected by centrifugation for 30 sec. at 14,000g, washed twice in PBS, resuspended in PBS, snap frozen into aliquots and stored at −80°C, or used directly for further processing.

For nucleolar isolation, freshly purified nuclei were treated according to Hacot et al. (43). In brief, purified nuclei were resuspended in nucleolar buffer (10 mM Tris, 10 mM NaCl, 2 mM MgCl_2_, pH 7.4 + Serva Protease Inhibitor mix HP (Serva Electrophoresis GmbH, Heidelberg, Germany)) containing 340 mM sucrose, and sonified on ice with a Branson Sonifier 250 (Branson Ultrasonics, Danbury, CT) until nuclear disruption was complete. Nucleoli were enriched by gradient centrifugation into nucleolar buffer containing 880 mM sucrose, and resuspended in 50 µl nucleolar buffer containing 340 mM sucrose, snap frozen, and stored in aliquots at −80°C.

The quality of the preparations and the organelle integrity was routinely analyzed by fluorescence staining with Hoechst 33342 (nuclei), or with the RNA-specific stain Sybr Green II (nucleoli).

To study SERF1a localization on isolated organelles, 10 µl of a nuclear or nucleolar preparation was incubated 5 min. with 10 µM Atto647N-SERF1a or Atto647N-K17E, centrifuged for 1 min. at 14,000g, and resuspended in 50 mM Tris, 150 mM NaCl, pH 7.4. To study RNA localization in isolated organelles, 10 µl of a nuclear or nucleolar preparation was incubated 5 min. with 1µM Cy3-R21(-) or Cy5-R21(+), respectively, in the abence or presence of 10 µM SERF1a, centrifuged for 1 min. at 14,000g, and resuspended in 50 mM Tris, 150 mM NaCl, pH 7.4. Fluorescence microscopy images were recorded on a Zeiss Axio Observer Z1 inverted microscope (Zeiss, Jena, Germany).

### eGFP-directed nuclear pull-down, MS analysis, functional classification and clustering

HeLa cells from three nearly confluent 75 cm^2^ flasks (Greiner Bio-One, Kremsmünster, Austria) overexpressing eGFP-SERF1a (Fig. S4B) were washed once with PBS, and combined after tryptic digestion. All further steps were performed on ice or at 4°C. The pellet was washed twice with PBS, and nuclei were isolated as descried for SH-SY5Y cells in the section above. Freshly isolated nuclei were used to prepare nuclear extracts by enzymatic lysis for 90 min. at 4°C with Active Motif enzymatic shearing cocktail in 100 µl Complete Digestion Buffer (both from Active Motif), and centrifuged for 10 min. at 14,000g as described in the manufacturer’s instructions (www.activemotif.com/catalog/25/nuclear-complex-co-ip-kit). The sample was diluted to 1 ml by the addition of 50 mM Tris, 50 mM NaCl pH7.4 + Serva Protease Inhibitor mix HP (Serva Electrophoresis GmbH). RNAse was omitted in order to preserve the integrity of RNA-dependent interaction clusters during the pull-down procedure. 30 µl pre-washed anti-GFP agarose beads (MBL, Woburn, MA) were added to the supernatant and incubated overnight at 4°C on a rotary shaker. The sample was loaded on a 0.4 ml Pierce microspin column (Thermo Fisher Scientific) centrifuged for 30 sec. at 14,000g, and washed two times with 50 mM Tris, 150 mM NaCl, 1% Igepal, pH 7.4 and one time with 50 mM Tris, 150 mM NaCl, pH 7.4 Bound proteins were eluted with 50 mM Tris, 1 M NaCl pH 7.4. The eluate was mixed with denaturing, reducing 4x Laemmli buffer and proteins were resolved on a 12% NuPAGE gel (Thermo Fisher Scientific). Proteins were visualized by silver staining. Bands of interest were excised, proteins reduced, alkylated and digested with trypsin. Resulting peptides were extracted, analysed by nanoLC-MS/MS and identified by MASCOT analysis (Matrix Science) as described by Gesslbauer et al. (44). Proteins were classified with respect to their molecular and biological function using the PANTHER classification system (version 14.1) (45). A Fisher overrepresentation test with Bonferroni correction was applied using default setup values.

Network clustering was performed by the STRING algorithm (version 11.0) (46) using experiments and databases as active interaction sources, and a minimum required interaction cut-off of 0.7 (high confidence).

### Sequence alignment

The SERF1a amino acid sequence was locally aligned with the EMBOSS Water alignment tools (47) against a collection of 289 non-canonical RNA binding motifs (branches) retrieved from the RBDmap database (10). Each branch with an identity of ≥ 6 residues was selected for further manual alignment.

### Alternative splicing assay

pMT-E1A was kindly provided by J. F. Caceres (MRC Human Genetics Unit, Western General Hospital, Edinburgh). HeLa cells were seeded in 12well plates (Corning Ing, Corning, NY) and transiently co-transfected in triplicates with *pMT-E1A* and *pEGFP-SERF1a*, SERF-K17E, or empty pEGFP control vehicle using jetPRIME transfection reagent (Polyplus Transfection, Illkirch-Graffenstaden, France) according to the manufacturer’s instructions. For each well, 0.5µg of each plasmid (corresponding to 1 µg total DNA), 100 μl jetPRIME Buffer and 2 μl jetPRIME transfection reagent were employed. Transfection efficiency was assessed by visual inspection under UV light and was approx. 90% in all cases. RNA was isolated 48h post transfection by phenol-chloroform-extraction, and subjected to reverse transcription using the Dynamo cDNA Synthesis Kit (Thermo Fisher Scientific). E1A alternatively spliced isoforms were amplified from 3 μl cDNA by PCR in 50 μl reactions with the KAPA HiFi PCR Kit (Peqlab/VWR, Darmstadt, Germany) using primers E1A_ Exon1_forward 5’-GTT TTC TCC TCC GAG CCG CTC CGA-3’ and E1A_Exon2_reverse 5’-CTC AGG CTC AGG TTC AGA CAC AGG-3’ at final concentrations of 500 nM. PCR conditions were 5 min of denaturation, 30 PCR-cycles (20s/98°C, 15s/70°C and 30s/72°C), 5 min 72°C. For the quantification of E1A spliced, 1 μl of each PCR product was subjected to Chip-based capillary electrophoresis with the Agilent 2100 Bioanalyzer (Agilent, Santa Clara, CA) and the proportional amount of each fragment to the whole yield of splicing products was calculated. One-way ANOVA F-tests were performed using R version 2.9.1 for comparing levels of spliced isoforms.

### Electrophoretic mobility shift assay

100 µM SERF1a was incubated with increasing amounts of R21(-) and samples were run on a 8% native-PAGE gels (Tris⋅HCl, pH 7.6). Due to net charge inversion of the SERF1a:RNA complex, one gel was run with inverted electrodes. Bands were visualized by Coomassie blue staining.

### Fluorescence anisotropy titrations

The fluorescence anisotropy of 200 nM Cy3-R21(-) or 200 nM Atto550-SERF1a in 50 mM bis-Tris, 20 mM NaCl, 3mM NaN_3_, pH 6.8, was measured at 25°C on a LB50 spectrofluorimeter (Perkin Elmer, Waltham, MA) equipped with excitation and emission polarizers-Ex/Em wavelength of Cy3-R21(-) was 550/570 nm. Ex/Em wavelength of Atto550-SERF1a was 554/576 nm. Slits widths were 5 nm and 10 nm for excitation and emission, respectively. The fluorescence anisotropy is defined as (48)

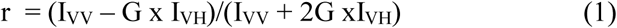

where I_VV_ is the fluorescence intensity recorded with excitation and emission polarizers in vertical positions, and I_VH_ is the fluorescence intensity recorded with the emission polarizer aligned in a horizontal position. The G factor is the ratio of the sensitivities of the detection system for vertically and horizontally polarized light G=I_HV_/I_HH_.

Binding curves were measured at low ionic strength (50 mM NaCl) to improve saturation. Cy3-R21(-) was titrated against increasing amounts of wtSERF1a, K17E, YDL085C-A, or MOAG-4 dialyzed against the same buffer. Atto550-labeled SERF11a was titrated against unlabeled R21(-), an analogous dsDNA 21-mer, heparin from porcine intestinal mucosa (MW_mean_ = 5,000 g/mol), tRNA from *S. cerevisiae* (type X-SA, MW ~ 25,000 g/mol; BioNumbers BNID 101177 (48)), or total RNA from *S. cerevisiae* (MW ~10^6^ g/mol by roughly approximating the composition of total RNA to 100% rRNA; BioNumbers BNID 105192 (49)). For each point, the anisotropy was recorded over 30 sec. and the mean r values for each measurement were used. Anisotropy changes were fitted by using the Levenberg-Marquardt algorithm to the equation *(2)*

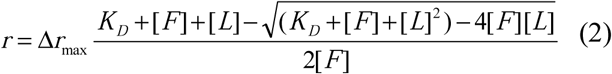

where r is the observed anisotropy, Δr_max_ is the maximal anisotropy change, and K_D_ is the dissociation constant. F indicates the fluorescence labelled binding partner, L is the ligand used for titrations.

### Isothermal titration calorimetry

Titrations were performed in 50 mM bis(2-hydroxyethyl)amino-tris(hydroxymethyl)methan (bis-Tris), 20 mM NaCl, 3mM NaN_3_, pH 6.8, at 25°C using a VP-ITC microcalorimeter (Microcal, Pittsburgh, PA). 1mM 21R(-) was titrated to a 30 µM SERF1a solution. The titration sequence consisted of 2 µl injections of 4 sec. each, with a spacing of 250 sec. between the injections, for a total of 20 injections. OriginLab software (Microcal) was used to fit the raw data to a 1∶1 binding model.

### RNA duplex annealing assay

The assay was performed as described by Rajkowitsch et al. (15), except for using a quartz cuvette instead of microplates. In brief, the complementary fluorophore-tagged RNA 21mers Cy5–21R(+) and Cy3–21R(-) were annealed at 37°C in 50 mM Tris⋅HCl (pH 7.5), 3 mM MgCl_2_, and 1 mM DTT in the absence and in the presence of 1µM SERF1a, and of aSyn when required. The time-resolved fluorescence emission of the FRET Cy3/Cy5 donor/acceptor pair was measured in a Cary Eclipse fluorescence spectrometer (Agilent, Santa Clara, CA, USA) with 10 nm excitation and emission slit widths. Emission was measured at 570 nm and 670 nm, respectively, upon donor excitation at 550 nm. The FRET efficiency E_FRET_ was calculated using the equation

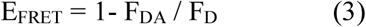

where F_DA_ and F_D_ are the fluorescence intensities of the donor in the presence and in the absence of the acceptor, respectively.

The reaction constant K_obs_ was calculated by the equation (15)

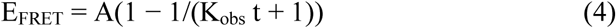

Where K_obs_ is the observed annealing reaction constant, and *A* is the maximum reaction amplitude.

### Generation, annealing and analysis of RNA-based complex coacervates

5 µl 20 µM Cy3-21R(-) or Cy5-21R(+) in 50 mM Tris⋅HCl, 1 mM MgCl_2_, pH 7.5 were mixed with 5µl of a 200 µM poly-L-arginine (polyR) hydrochloride (MW ~15,000-70,000 g/mol) solution in the same buffer (RNA:polyR = 1:10). The appearance of phase separated fluorescent liquid droplets of approximately 1 µm diameter was immediately observable. Complex coacervates were freshly prepared before each experiment.

For RNA duplex annealing under two-phase conditions, equal volumes of a Cy3-21R(-)/polyR and a Cy5-21R(+)/polyR complex coacervate preparation were mixed and incubated on a borosilicate coverslip for 5 min in the absence and in the presence of 10µM SERF1a.

Fluorescence images were recorded at room temperature under a Zeiss Observer Z1 inverted microscope equipped with appropriate fluorescent filter sets (Cy3: ex/em 549/562 nm; Cy5: ex/em 646/664 nm). Co-localization analysis was performed on three independent samples with ImageJ and the Colocalization Threshold plugin to obtain the Pearson correlation coefficient for each sample.

To study the incorporation of RNA-derived liquid droplets into nucleoli, 1 µl of isolated nucleoli was added to 9 µl Cy5-21R(+)/polyR complex coacervates and incubated for 5’ in the absence (-) or in the presence (+) of 10 µM SERF1a. Fluorescence images of each sample were recorded at room temperature under a Zeiss Observer Z1 inverted microscope. Nucleoli were counterstained with Sybr Green II RNA dye (ex/em filter 495/517 nm).

### NMR spectroscopy

NMR measurements were carried out at 280K on a Bruker Avance III 700-MHz spectrometer equipped with cryogenically cooled 5 mm TCI probe. 21R(-) at final concentrations of 12, 25, 49 and 96 µM was added to 50 µM uniformely ^15^N-labeled SERF1a in 50 mM bis-Tris, 20 mM NaCl, 3mM NaN_3_, pH 6.8. ^1^H,^15^N-HSQC spectra were recorded at each titration step. {^1^H}-^15^N heteronuclear NOE and HNCACB spectra of 200 µM uniformly ^13^C,^15^N-labeled SERF1a with and without 200µM 21R(-) were measured at 700 MHz as described in Ref. 26. Backbone resonance assignment of SERF1a was performed as previously described (4). Spectra were processed using NMRPipe (50) and analyzed with Sparky ((T. D. Goddard and D. G. Kneller, SPARKY 3, University of California, San Francisco).

### FRET-based competition assay

50 nM Atto550-SERF1a was mixed with equimolar amounts of Cy5-R21(+) and FRET of the Atto550/Cy5 donor/acceptor pair was measured in a Cary Eclipse fluorescence spectrometer with 10 nm excitation and emission slit widths. Emission was measured at 576 nm and 670 nm, respectively, upon donor excitation at 554 nm. Changes of the FRET signal were measured upon the addition of C-terminal aSyn peptide. The normalized FRET emission ratio was normalized, and the resulting curve was fitted for one–site competitive binding using the analysis program Origin 8.0 (Microcal).

## ACKNOWLEDGEMENTS

We thank Ute Schäfer (Medical University Graz) for mouse brain samples.

## COMPETING INTERESTS

The authors declare no competing financial interest.

## AUTHOR CONTRIBUTIONS STATEMENT

N.H.M. was repsonsible for NMR measurements and data processing. H. D. and J. G. planned and carried out alternative splicing assays. D.A.M. and M.G. carried out protein mutagenesis. A.L. performed protein purification. C.T. prepared cellular and brain tissue sections. C.T. and N.H.M. performed immunofluorescence analysis. R.P. performed histological analyses of hippocampal mouse tissue. H.D. and S. F. F. carried out living cell fluorescence microscopy. S.F.F performed pull-down analysis, ITC, fluorescence-based *in vitro* experiments, qPCR, and bioinformatic analyses. T. M. F. and S. F. F. carried out co-localisation analyses. B.G. and J.A. performed MS-analysis. E. A. N. provided MOAG-4 protein. K.Z. provided technical NMR advice. A.J.K. provided access to cell culture and MS-facilities. S.F.F redacted the manuscript.

