## Supplemental Data for "Structural fuzziness of the RNA-organizing protein SERF1a determines a toxic gain-of-interaction"

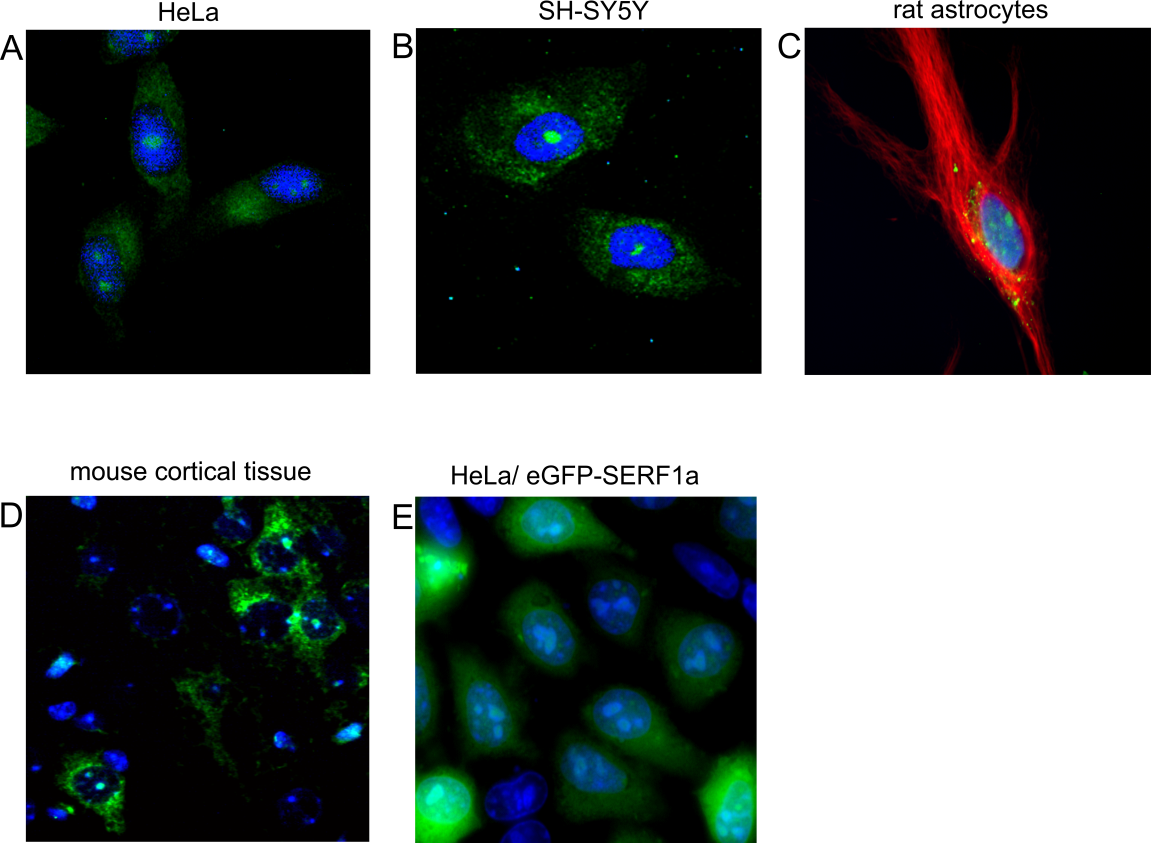

FIGURE S1: (A-D) IF-images showing the intracellular distribution of constitutively expressed SERF1a in different fixed cell type preparations. Nuclei are stained with DAPI (blue). Rat astrocytes in mixed glial preparations (C) are stained with glial fibrillary acid protein as a specific marker (red). (E) Distribution of transiently overexpressed eGFP-SERF1a (green) in living HeLa cells. Nuclei are stained with Hoechst 33342.

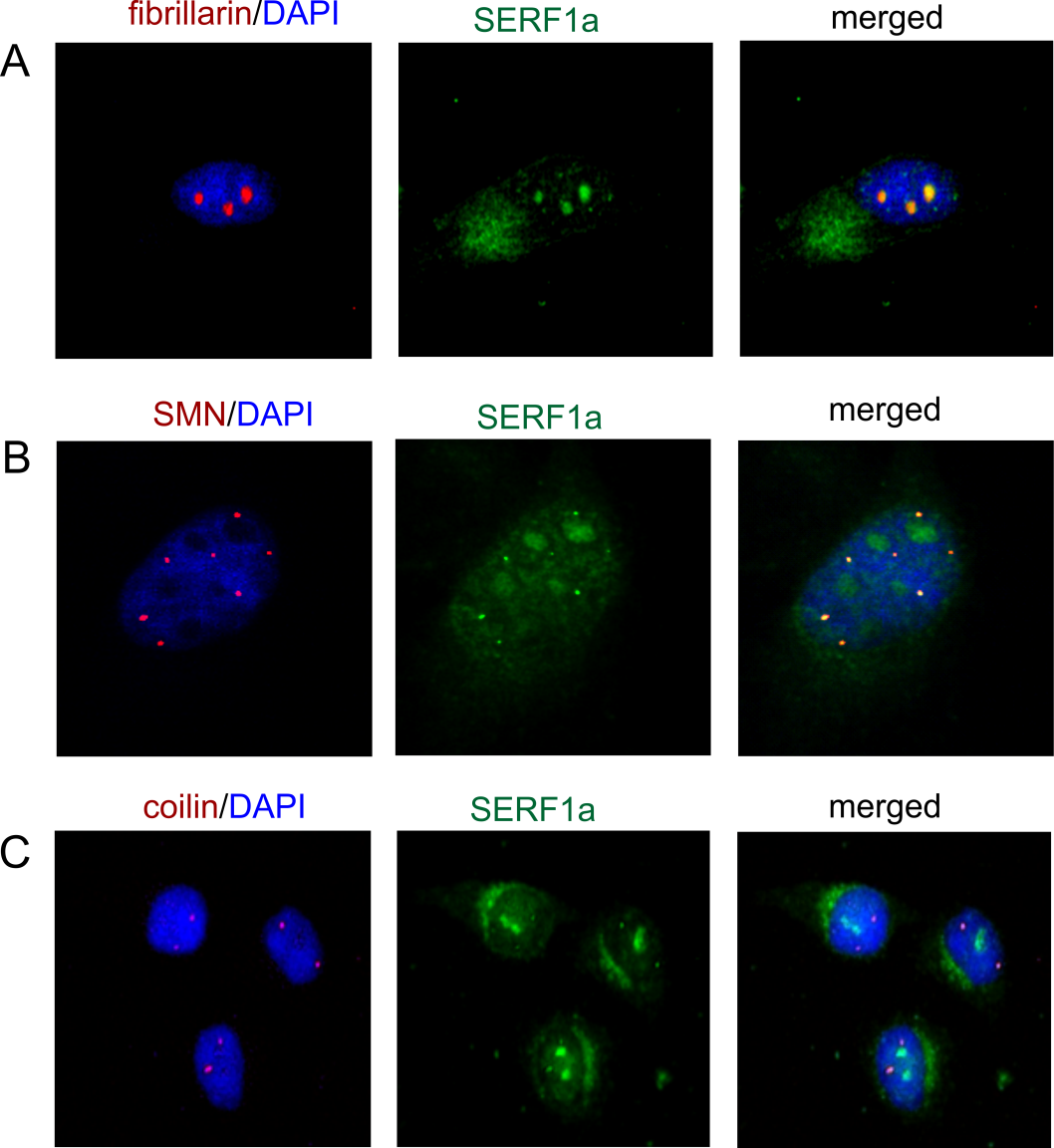

FIGURE S2: intranuclear localization of endogenous SERF1a in fixed HeLa cells. (A) SERF1a (green) and nucleolar marker fibrillarin (red) (B) SERF1a (green) and marker of nuclear gems SMN (red). (C) SERF1a (green) and Cajal Body marker coilin (red). Nuclei are stained with DAPI (blue).

*TIF-IA cKO mouse model*

To analyze the subcellular localization of SERF1a protein in neurons, we performed immunohistochemistry by a SERF1a specific antibody on paraffin sections of adult mouse brains. In hippocampal neurons SERF1a is localized in both cytoplasm and nucleoli (Figure 1D, upper panel), confirming the pattern observed in other cell lines.

To investigate whether SERF1a localization is linked to nucleolar activity, we analyzed hippocampal neurons of TIF-IA^CaMKCreERT2^ mutant mice (TIF-IA cKO) (40). These mice are characterized by inhibition of rRNA transcription and perturbed nucleolar integrity 4 weeks after induction of TIF-IA genetic ablation by tamoxifen treatment in adult mice (39, 40). In this model of nucleolar stress, SERF1a was absent in the nucleoli, while still localized in the cytoplasm (Figure 1D, lower panel). The analysis of SERF1a expression by RT-qPCR in controls and TIF-IA cKO mice at the same stage (Fig. S3) indicates that SERF1a levels are similar in control and mutant mice (unpaired t-test, p=0.09). These results suggest that the different SERF1a immunoreactivity is independent of changes in SERF1a mRNA.

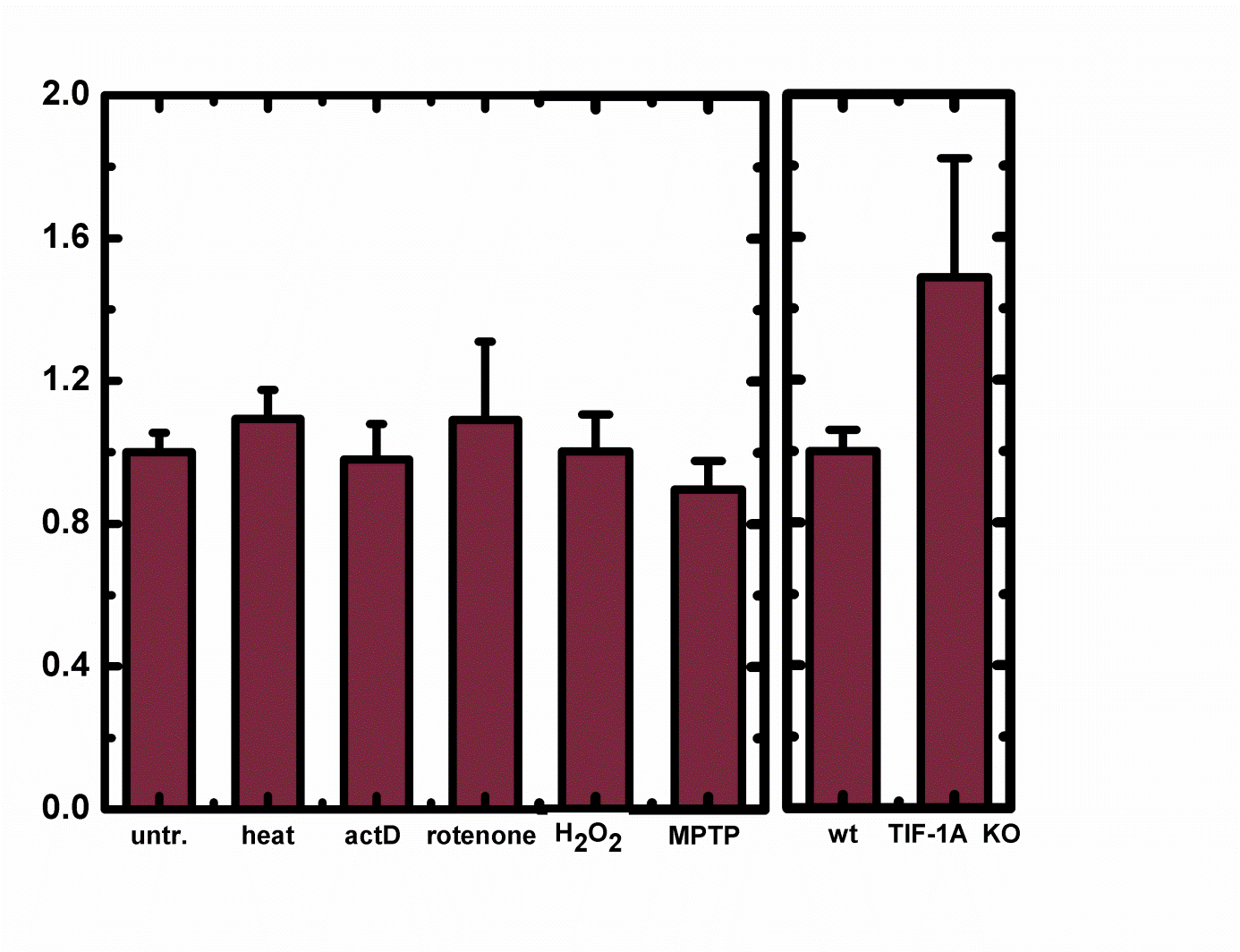

FIGURE S3: qPCR analysis of *SERF1a* gene expression under different stress conditions. Left panel: SERF1a mRNA levels of SH-SY5Y cells after exposure to heat, actinomycin D, rotenone, hydrogen peroxide, or MPTP. (related to Fig. 1A-C). Error bars represent SEM (n=3). Right panel: SERF1a mRNA levels in control and TIF-IA cKO mice 4 weeks after tamoxifen injection. Error bars represent SEM (n=5). GAPDH was used as internal control. Expression changes were calculated as a fold change vs. mean of control samples.

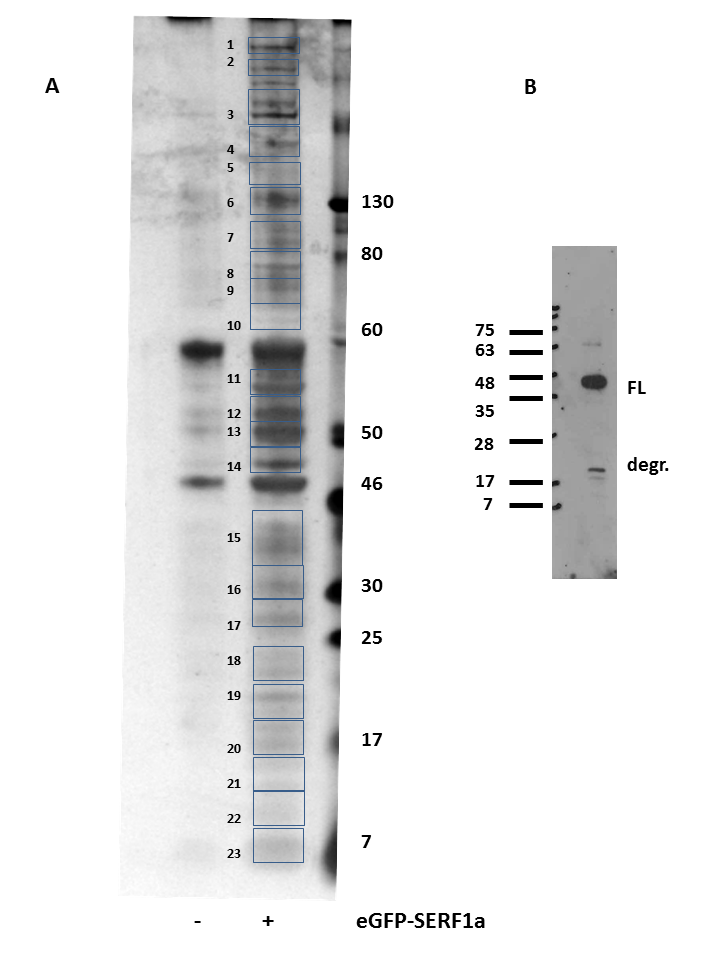

FIGURE S4: (A) SDS-PAGE and silver staining analysis of proteins captured by anti-GFP agarose out of (-) untransfected HeLa control cells and (+) cells overexpressing eGFP-SERF1a. (B) SERF1a-directed immunoblotting of cell lysates overexpressiong eGFP-SERF1a. FL: full length eGFP-SERF1a (approx.. 35 kDa) ; degr. : degradation products

*Cellular E1a minigene splicing model*

We tested for a trans-regulatory effect of SERF1a on RNA splicing in a living cell exon splicing model (S1). We found that the overexpression of eGFP-SERF1a in HeLa cells led to splice-site variations in the transcription of the adenoviral E1A reporter minigene. By qPCR analysis of E1A splicing variants in eGFP-SERF1a transfected cells (Fig. 5B), we could observe an increase of 10S mRNA levels by 7.6% (from 25.6% to 33.2%), in concomitance to a decrease of 11S mRNA by 3.3% (from 5.3% to 2.0%), and of 12S mRNA by 5.9 % (from 7.7% to 1.8%). Less significant changes were observable for 9S, 11S, and 13S isoforms. These results suggest that SERF1a favours the use of different splice sites respectively to an unfused eGFP control. Such effect was far less accentuated for the RNA-defective binding mutant K17E. Respectively to an unfused eGFP control, the expression of 10S transcripts was increased only by 2.7% (from 25.6% to 28.3%). Concomitantly, 11S mRNA and 12mRNA levels decreased by only 0.8 % (from 5.3% to 4.5%) and 2.1% (from 7.7% to 5.6%), respectively. These findings suggest that SERF1a exhibits a trans-splicing-associated function deriving from a direct interaction with RNA.

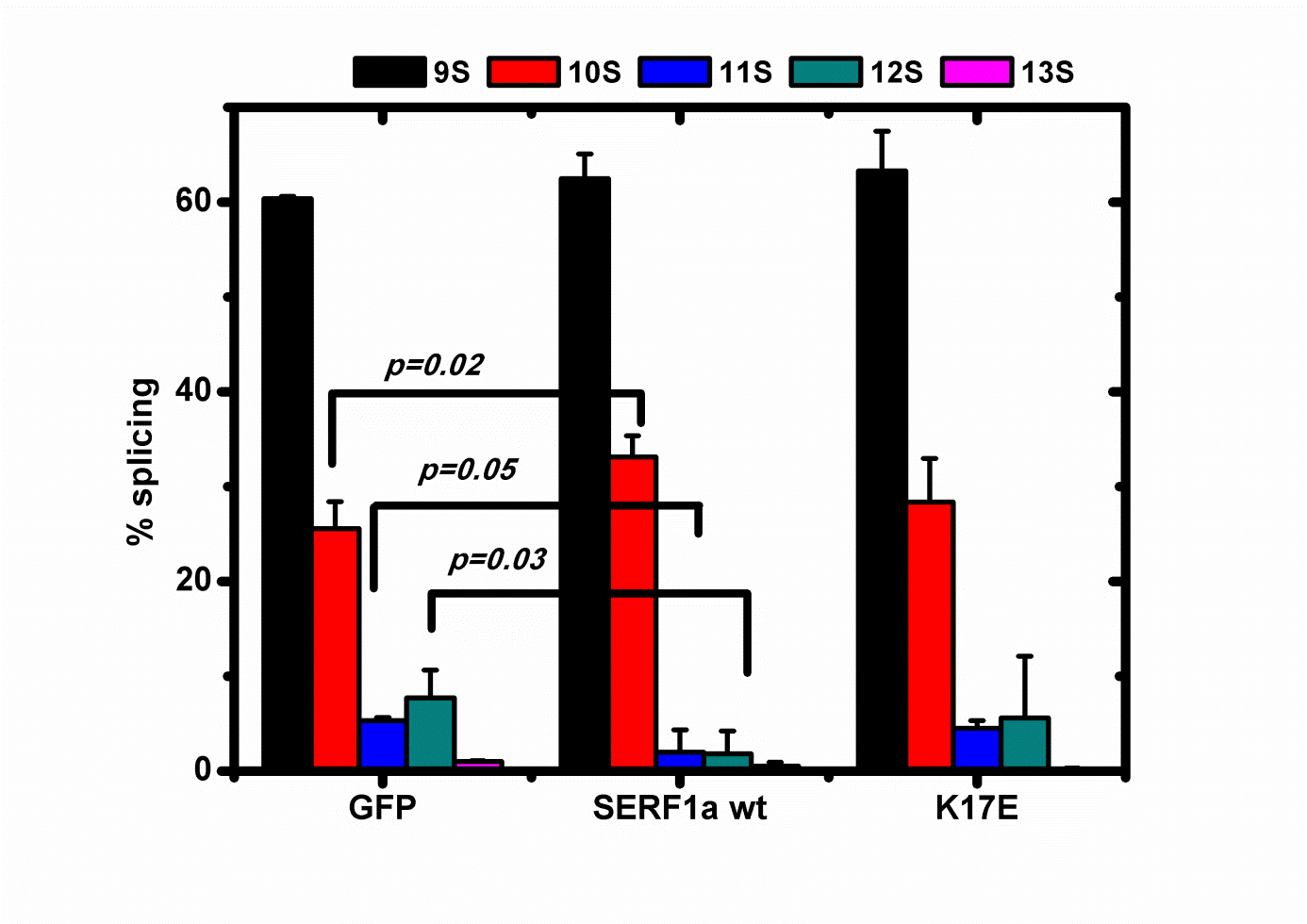

FIGURE S5: Regulatory effects of SERF1a on RNA splicing. E1A minigene living cell alternative splicing assay showing splicing variations of major mRNA transcripts upon the overexpression of eGFP-SERF1a, eGFP-K17E, and eGFP control vehicle (n=3).

*Variations in pre-rRNA levels*

The functional relevance of SERF1a in the nucleolus was assessed by cellular 47S pre-rRNA levels as a typical signature of ribosome biogenesis (S2, S3), and thus of nucleolar activity. Variations in pre-rRNA levels were measured by qPCR with an oligonucleotide targeting the A0 site within the 5’ externally transcribed region (5′ETS) of 47S (S2, S3). We observed that A0 levels varied upon SERF1a downregulation or overexpression (Fig. S4). Silencing of SERF1a led to an approximately fourfold amplification increase. A return towards initial levels of 47S pre-rRNA could be observed by the subsequent overexpression of eGFP-SERF1a, while the RNA-binding defective SERF1a mutant K17E failed to provide a similarly pronounced effect. These results suggest that SERF1a can influence rRNA biogenesis via a direct interaction with RNA.

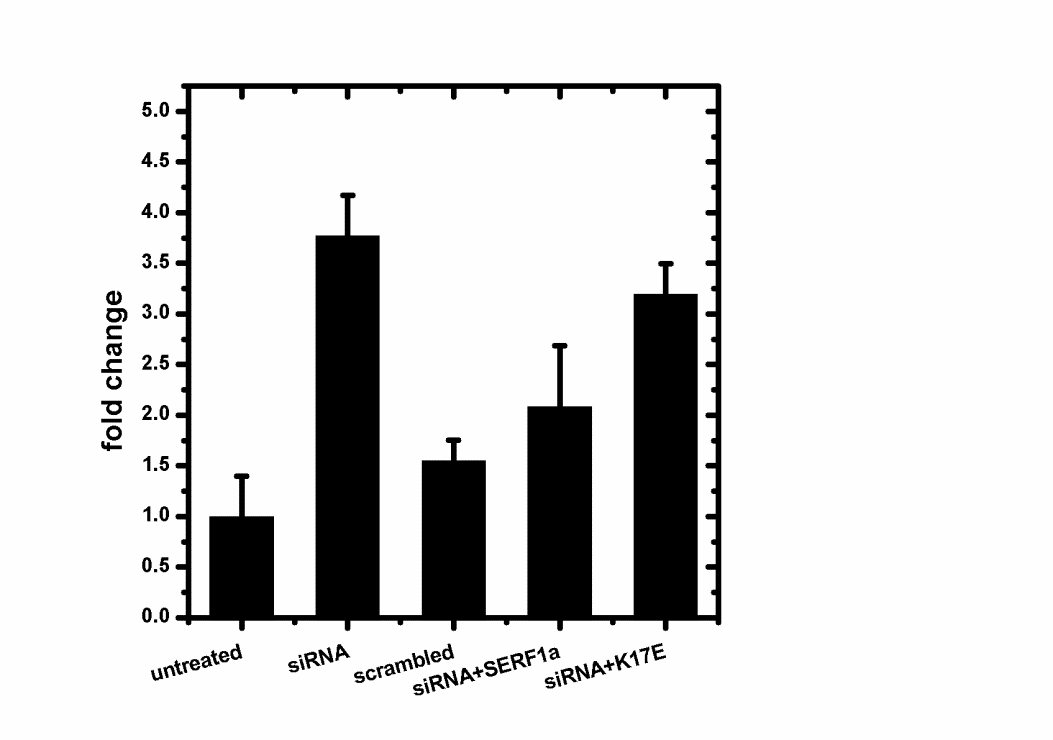

FIGURE S6: qPCR analysis of 47S pre-rRNA (A0) levels in SH-SY5Y cells. GAPDH was used as internal control. Expression changes were calculated as a fold change vs. mean of control samples (n=3).

**A**

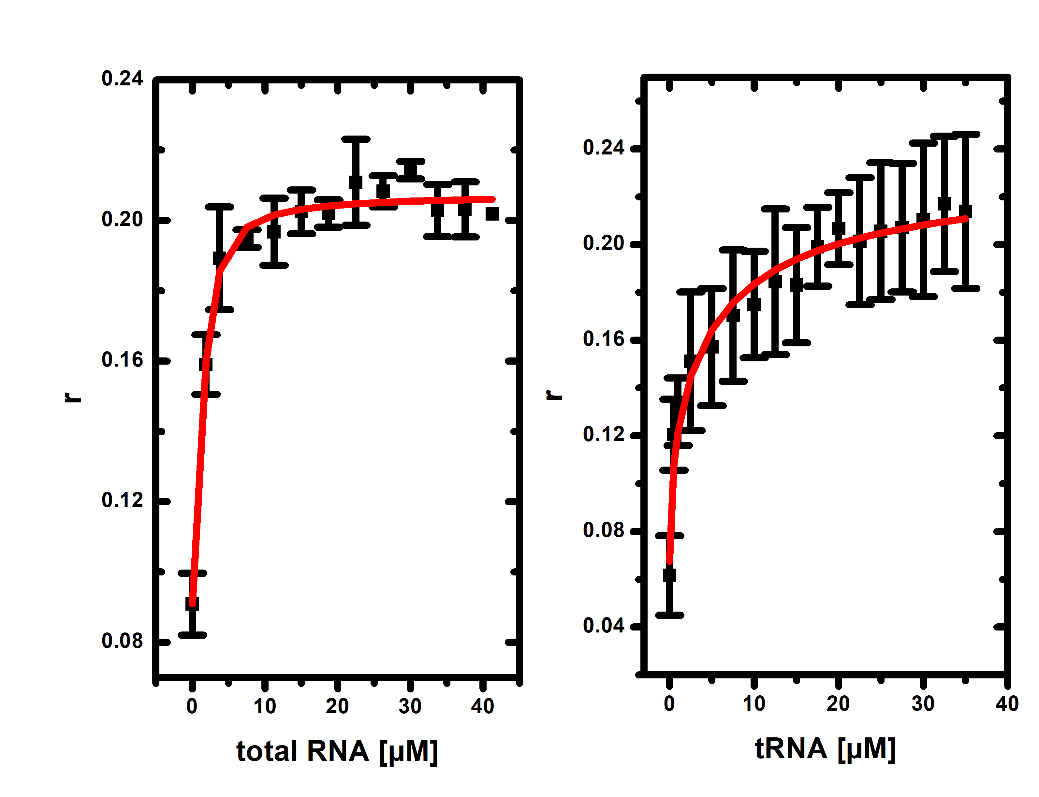

**B**

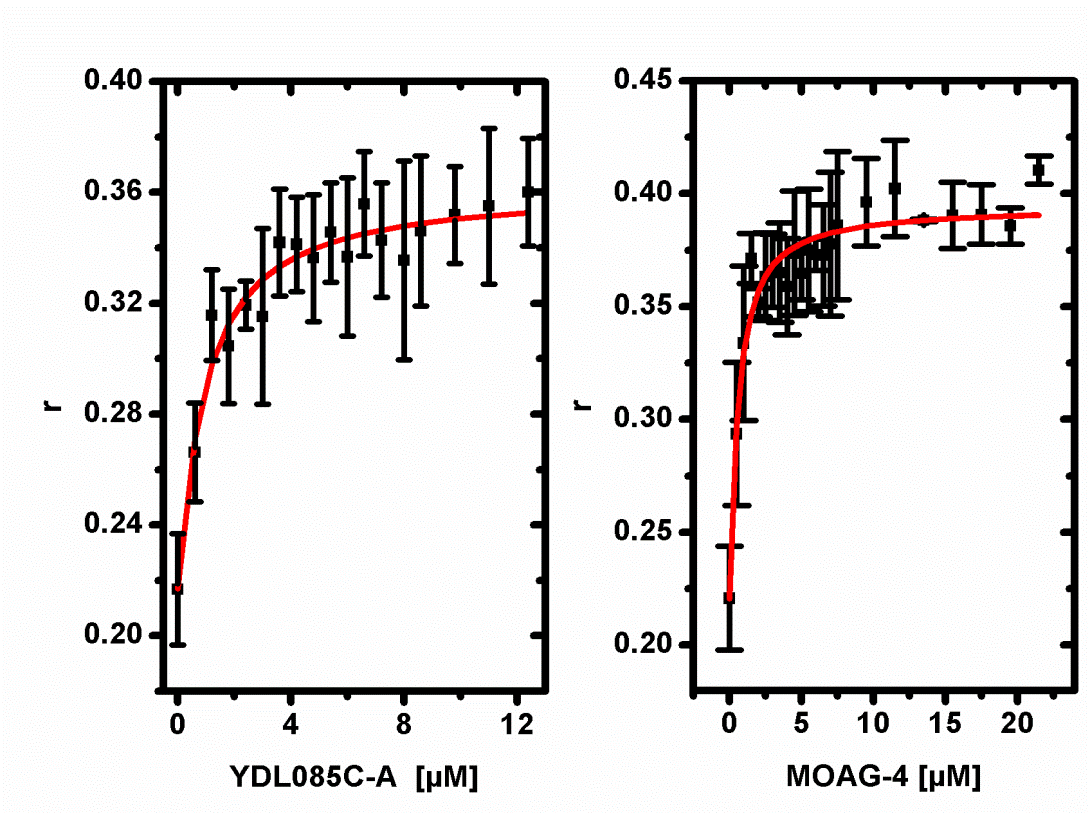

FIGURE S7: (A) Fluorescence anisotropy saturation curves obtained by titration of Atto550-SERF1a with total RNA from *S. cerevisiae* (left panel, or with tRNA from *E. coli* (right panel). (B) Fluorescence anisotropy saturation curves obtained by titration of Cy3-R21(-) with the SERF1a orthologues YDL085C-A from *S. cerevisiae* (left panel) or MOAG-4 from *C. elegans*

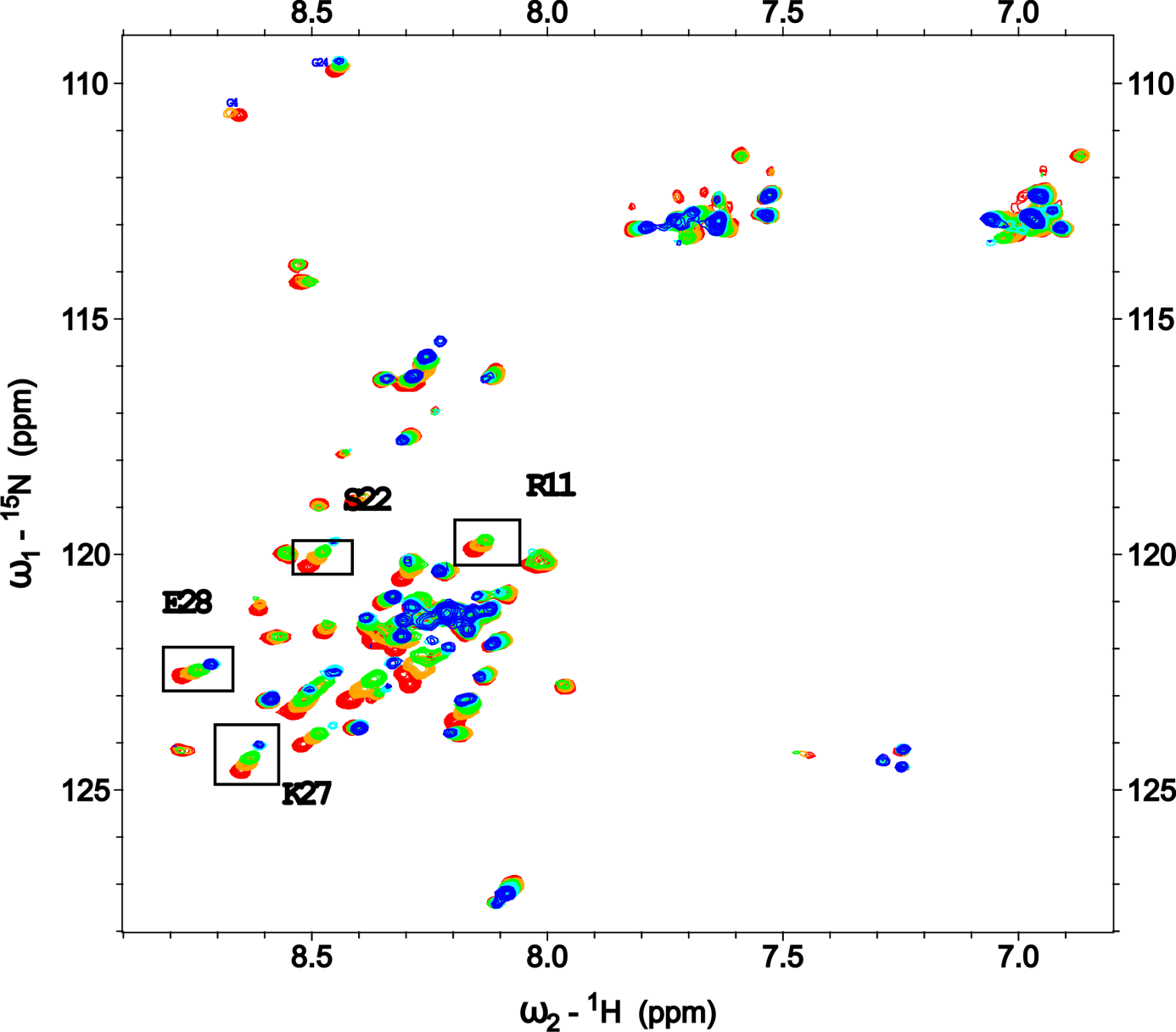

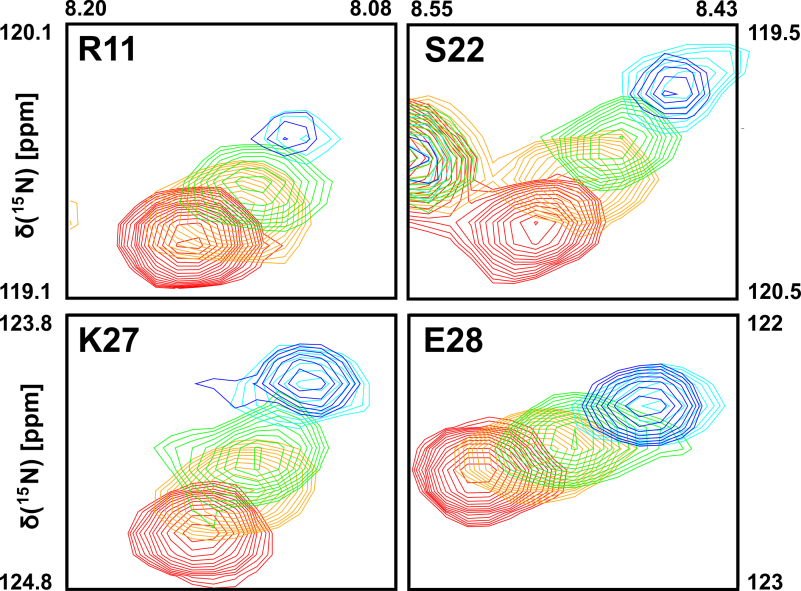

FIGURE S8: Overlay of SERF1a ^1^H,^15^N-HSQC NMR spectra titrated with increasing ratios of R21(-). Four well-resolved SERF1a chemical signals (residues R11, S22 and K27 and E28) shifting upon titration with RNA are magnified in detail.

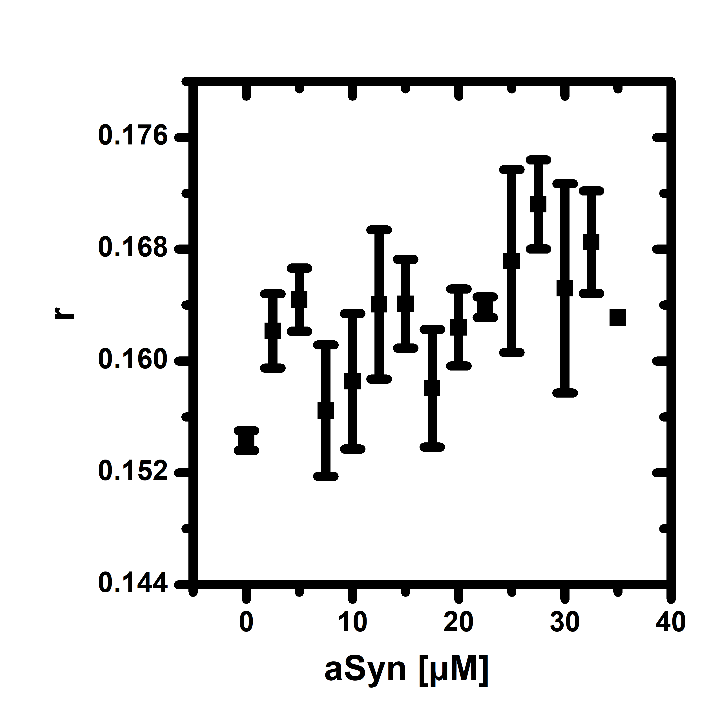

FIGURE S9: Fluorescence anisotropy saturation curve obtained by titration of Cy3-R21(-) with aSyn.

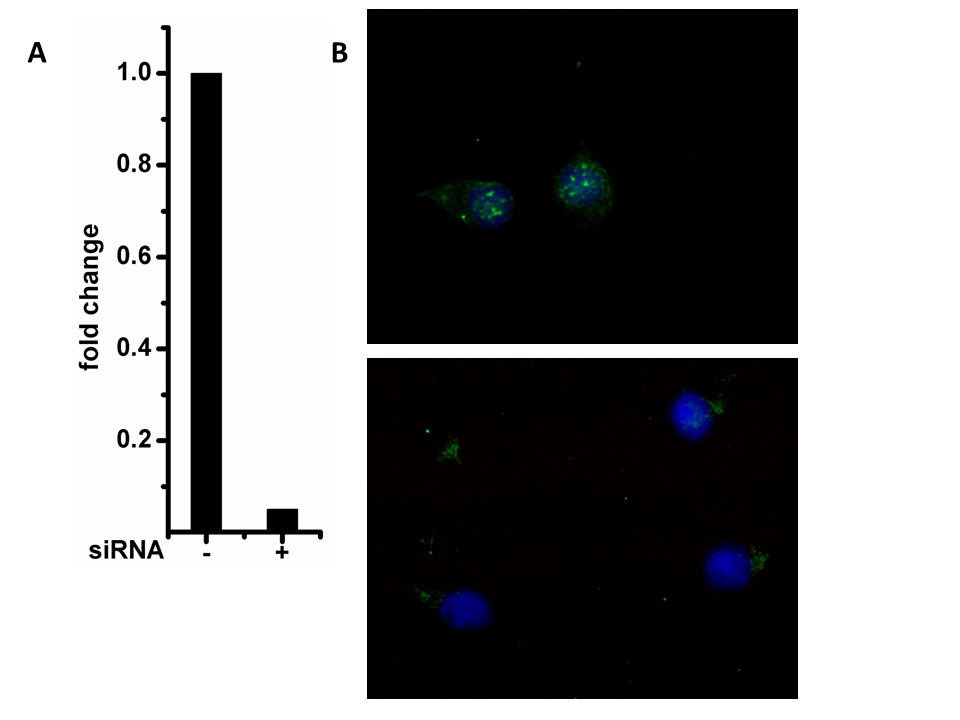

FIGURE S10: silencing of SERF1a in SH-SY5Y cells. (A) qPCR analysis of *SERF1a* gene expression levels. GAPDH was used as internal control. (B) IF analysis of unsilenced (upper panel) and silenced (lower panel) cells. SERF1a is green. Nuclei are counterstained with DAPI (blue).

TABLE S1: Pull-down database (see separate file TableS1.xls)

TABLE S2: *qPCR primers*

| GAPDH | 5’ ATGTTCGTCATGGGTGTGAA  3’ GTCTTCTGGGTGGCAGTGAT |
| --- | --- |
| SERF1, mouse | FM1_Serf1 (8019015181-10/0) AAACATGAAAAAGACCCAGG (Sigma Aldrich)  RM1_Serf1 (8019015181-10/1) ATGATCTCTGAATCCCTCTG (Sigma Aldrich) |
| SERF1a, human | FH1_SERF1b (8015430739-100001) AGGAAAGAGGATAGCTTGAC (Sigma Aldrich)  RH1_Serf1b (8015430739-100002) GACTGTCCACTCTACAAATC (Sigma Aldrich) |
| 5’ETS(A0) | 5’ GAACGGTGGTGTGTCGTTC (5)  3’ GCGTCTCGTCTCGTCTCACT |

TABLE S3: list of used plasmids

| *Plasmid name* | *Product* | *Reference* |
| --- | --- | --- |
| *pJ414/ YDL085C-A* | YDL085C-A | commercial synthetic plasmid |
| *pEGFP/SERF1a* | eGFP-SERF1a | 4 |
| *pEGFP/K17E* | eGFP-K17E | this paper |
| *pcDNA/mCherry-SERF1a* | mCherry-SERF1a | 1 |
| *pcDNA/mCherry-K17E* | mCherry-K17E | 4 |
| *EGFP-alphasynuclein-WT* | eGFP-aSyn | gift from David Rubinsztein (Addgene #40822); S4 |
| *pRECEIVER-B02/SERF1A* | SERF1a | 2 |
| *pRECEIVER-B02/SERF1AK17E* | K17E | 4 |
| *pRSETB/SNCA* | aSyn | S5 |
| *pMT-E1A* | E1A minigene | gift from J.F. Caceres |
